## Supplementary Information for "Benchmarking and behavioral characterization of LLM agents for protein design"

### Appendix A Methods

#### A.1 Task Design and Taxonomy

The resulting $2 \times 5$ matrix yields 10 possible taxonomy cells, of which 9 are populated. The single empty cell is biologically justified: binder × redesign = 0 because *de novo* binder design via RFdiffusion is the current state-of-the-art and no established redesign workflows exist for miniprotein binders. One cell is nearly empty: fluorescent protein × de novo contains only 1 task, reflecting that GFP-family optimization via directed evolution is the dominant approach and de novo fluorescent protein design remains nascent. The distribution imbalance (47:29 de novo:redesign) reflects the current protein design landscape, where generative methods dominate binder and scaffold design while redesign approaches dominate enzyme and fluorescent protein optimization.

Each task is specified as a JSON object following a Pydantic-validated `DesignTask` schema (`biodesignbench/tasks/schema.py`). The schema includes fields for task identification (`task_id`, `description`), a free-form `task_type` string used for downstream taxonomy lookup, design `constraints` (`time_limit_minutes`, `knowledge_cutoff`, `network_allowed`), `metadata` (difficulty, source, DOI, expected tools, tags), `target` specification (input PDB, chain selectors, binding-site residues), `design_constraints` (length range, mutation budget, required residues), and an `evaluation` block (method, expected metrics, ground-truth path). The two-axis taxonomic classification (`DesignApproach`, `MolecularSubject`) is not a stored field but is derived deterministically from the task ID prefix at evaluation time via `taxonomy.get_category()`, which matches against the canonical regex `^([a-z]{2})-([a-z]{2,3})-(\d{3})$` (e.g., `dn.bnd_001` → (DE\_NOVO, BINDER)). Sensitive metadata fields (`source`, `doi`, `tools_expected`, `tags`, `test_file`) and the `evaluation.ground_truth` pointer are stripped before agent execution by the data sanitizer (Section A.6).

Tasks are assigned one of three difficulty levels using a deterministic three-tier fallback system. First, if a manual difficulty annotation exists in the task metadata, it is used directly. Second, if the task’s `design_function` has a predefined difficulty mapping (e.g., `BACKBONE_GENERATION` → difficulty 3, `SEQUENCE_DESIGN` → difficulty 1), that mapping is applied. Third, as a final fallback, difficulty is inferred from the number of required design functions: tasks requiring $\leq 2$ functions are assigned difficulty 1 (easy; 21 tasks), tasks requiring 3–4 functions are assigned difficulty 2 (medium; 28 tasks), and tasks requiring $\geq 5$ functions are assigned difficulty 3 (hard; 27 tasks). This produces a balanced distribution across the three levels.

Each task is presented to the agent as a natural language prompt that specifies the design objective, input structure, constraints, and expected outputs. The prompt is generated programmatically from the task JSON and includes an anti-contamination instruction (Section A.6) that explicitly prohibits the agent from searching for published solutions. A representative prompt reads: “Design a nanobody that binds the RBD domain of SARS-CoV-2 spike protein. Input PDB: [path]. Target chains: A. Design chains: B. Constraints: length 110–130 residues. Generate at least 3 diverse designs with predicted structures. Do NOT search for or reference any published papers, DOIs, or known solutions for this task.”

To decouple evaluation from specific tool names, we define 10 abstract design functions that represent the fundamental computational operations in protein design: `BACKBONE_GENERATION` (generating new protein backbone coordinates), `SEQUENCE_DESIGN` (designing amino acid sequences for a given backbone), `STRUCTURE_PREDICTION` (predicting 3D structure from sequence), `COMPLEX_PREDICTION` (predicting multi-chain complex structures), `INTERFACE_ANALYSIS` (analyzing protein-protein or protein-ligand interfaces), `STABILITY_SCORING` (assessing thermodynamic stability), `ENERGY_MINIMIZATION` (relaxing structures to local energy minima), `HOTSPOT_IDENTIFICATION` (identifying critical binding residues), `SEQUENCE_SCORING` (evaluating sequence fitness), and `PHYSICS_VALIDATION` (validating biophysical properties). Each MCP tool maps to one or more design functions (Table S1), and the approach scoring component (Section A.3) evaluates which design functions were executed rather than which specific tools were invoked.

**Table S1** Mapping between MCP tools and abstract design functions. Each tool maps to one or more design functions; approach scoring credits agents for executing the correct functions regardless of which specific tool was invoked. UG = available in both unguided and guided modes; GD = guided-only (composite multi-step pipelines hidden in unguided mode so agents must orchestrate atomic tools themselves).

| MCP Tool | Design Function(s) | Mode |
| --- | --- | --- |
| *Composite pipelines (guided-only)* |  |  |
| `design_binder` | `BACKBONE_GENERATION`, `SEQUENCE_DESIGN`, `STRUCTURE_PREDICTION` | GD |
| `design_fold` | `BACKBONE_GENERATION`, `SEQUENCE_DESIGN`, `STRUCTURE_PREDICTION` | GD |
| `optimize_sequence` | `SEQUENCE_DESIGN` | GD |
| *Atomic tools (available in both modes)* |  |  |
| `generate_backbone` | `BACKBONE_GENERATION` | UG |
| `design_sequence` | `SEQUENCE_DESIGN` | UG |
| `rosetta_design` | `SEQUENCE_DESIGN` | UG |
| `predict_structure` | `STRUCTURE_PREDICTION` | UG |
| `predict_structure_boltz` | `STRUCTURE_PREDICTION` | UG |
| `validate_design` | `STRUCTURE_PREDICTION` | UG |
| `predict_complex` | `COMPLEX_PREDICTION`, `STRUCTURE_PREDICTION` | UG |
| `predict_affinity_boltz` | `COMPLEX_PREDICTION`, `INTERFACE_ANALYSIS` | UG |
| `analyze_interface` | `INTERFACE_ANALYSIS` | UG |
| `rosetta_interface_score` | `INTERFACE_ANALYSIS` | UG |
| `rosetta_score` | `PHYSICS_VALIDATION` | UG |
| `score_stability` | `STABILITY_SCORING` | UG |
| `energy_minimize` | `ENERGY_MINIMIZATION` | UG |
| `rosetta_relax` | `ENERGY_MINIMIZATION` | UG |
| `suggest_hotspots` | `HOTSPOT_IDENTIFICATION` | UG |
| `get_design_status` | (utility) | UG |

### A.2 MCP Tool Framework

BioDesignBench provides agents with access to 19 protein design tools integrated through the Model Context Protocol (MCP) [1]. These tools are organized into three groups based on their underlying computational engine. The first group comprises 13 deep-learning-based tools: `design_binder` (target-conditioned RFdiffusion → ProteinMPNN → ESMFold pipeline for *de novo* binder generation), `design_fold` (unconditional RFdiffusion → ProteinMPNN → AlphaFold2 pipeline for *de novo* fold design), `generate_backbone` (unconditional RFdiffusion backbone generation), `design_sequence` (single-pass ProteinMPNN sequence design for a given backbone), `optimize_sequence` (ESM2-guided iterative mutation scanning for sequence optimization), `predict_structure` (AlphaFold2 / ESMFold monomer structure prediction), `predict_complex` (AlphaFold2-Multimer complex structure prediction), `validate_design` (sequence-input structure prediction with confidence aggregation), `analyze_interface` (binding interface analysis with buried surface area, hydrogen bonds, and contact maps), `score_stability` (ESM2 pseudo-log-likelihood stability scoring), `energy_minimize` (OpenMM energy minimization), `suggest_hotspots` (binding hotspot identification via computational alanine scanning), and `get_design_status` (asynchronous job status polling). The second group comprises 4 PyRosetta-based tools: `rosetta_score` (Rosetta energy function scoring), `rosetta_relax` (Rosetta FastRelax protocol), `rosetta_interface_score` (Rosetta interface energy decomposition with per-residue contributions), and `rosetta_design` (single-pass Rosetta PackRotamers+Min fixed-backbone design). The third group comprises 2 Boltz-based tools: `predict_structure_boltz` (Boltz-2 structure prediction) and `predict_affinity_boltz` (Boltz-2 binding affinity prediction).

Each tool is exposed to the agent as an MCP tool definition comprising a tool name, a natural language description of its functionality and intended use cases, and a JSON Schema specifying its input parameters and output format. Tool descriptions are designed to be informative without revealing optimal pipeline strategies; for example, the `design_binder` tool description states that it generates *de novo* binder proteins using RFDiffusion and explains the required inputs (target PDB, hotspot residues, number of designs) and expected outputs (designed PDB structures with confidence metrics), but does not specify when in a design pipeline it should be invoked relative to other tools. The MCP server communicates with agents via the stdio transport protocol, where tool invocations and results are exchanged as JSON-RPC messages over standard input/output streams.

To investigate how tool presentation affects agent performance, we evaluate all LLMs under two distinct MCP tool modes that differ in whether multi-step composite pipelines are exposed alongside their atomic constituents (Table S1). In unguided mode, agents receive 16 atomic tools listed individually with minimal descriptions — each tool is described in isolation without reference to other tools or pipeline strategies, and the system prompt provides no workflow hints. The 3 composite tools (`design_binder`, `design_fold`, `optimize_sequence`), which internally chain RFDiffusion, ProteinMPNN, and structure prediction or run iterative ESM2 mutation scanning, are withheld so the agent must discover and compose these multi-step workflows from first principles using the underlying atomic tools (`generate_backbone`, `design_sequence`, `predict_structure`, ...). In guided mode, agents receive all 19 tools with enhanced system-prompt guidance that includes four canonical workflow patterns (de novo binder design, sequence optimization, conformational design, and complex engineering), each spelling out the recommended atomic-tool sequence and noting when a composite tool can substitute for several atomic steps. This mode simulates a realistic deployment scenario where a protein-design practitioner has configured their MCP server with workflow-aware documentation.

The mode switching is implemented at the BioDesignBench tool provider level (`biodesignbench/tools/protein_design_provider.py`) through a `ToolMode` enum that controls (i) which tool definitions are surfaced to the agent SDK (composites are filtered out in `benchmark` mode via the `COMPOSITE_TOOLS` frozen set) and (ii) which system-prompt guidance string is prepended (`_BENCHMARK_GUIDANCE` vs. `_USER_GUIDANCE`). Both modes invoke identical underlying computational backends in the MCP server; only the tool metadata and system-prompt context visible to the agent differ.

Each MCP tool is mapped to one or more of the 10 abstract design functions defined in Section A.1 via the `TOOL_TO_FUNCTION` dictionary (`biodesignbench/eval/metrics/approach.py`). For example, `design_binder` and `design_fold` each map to `{BACKBONE_GENERATION, SEQUENCE_DESIGN, STRUCTURE_PREDICTION}` because they internally chain `RFDiffusion` → `ProteinMPNN` → structure prediction; `predict_structure` maps to `{STRUCTURE_PREDICTION}`; `predict_complex` to `{COMPLEX_PREDICTION, STRUCTURE_PREDICTION}`; and `predict_affinity_boltz` to `{COMPLEX_PREDICTION, INTERFACE_ANALYSIS}`. This mapping enables the approach scoring component to credit agents for executing the correct computational operations regardless of which specific tool they chose — an agent that uses `rosetta_interface_score` receives the same `INTERFACE_ANALYSIS` credit as one that uses `analyze_interface`.

All tool executions occur within a Docker container that provides a reproducible computational environment with pre-installed dependencies: PyRosetta, ColabFold/AlphaFold2, RFDiffusion, ProteinMPNN, ESM-2, and Boltz-2. The container is configured with GPU passthrough for GPU-accelerated structure prediction and backbone generation, and CPU limits to prevent the MCP server from consuming all available cores. Network access within the container is disabled to enforce the data contamination defense (Section A.6). Each agent session receives a fresh container instance with an isolated filesystem, ensuring that outputs from one task cannot influence subsequent tasks.

#### A.3 Scoring Rubric

Agent outputs are evaluated using a six-component, 100-point scoring rubric that combines algorithmic metrics (72 points) with rubric-based LLM judge assessment (28 points). The components are: Approach (20 points; 10 algorithmic + 10 LLM), Orchestration (15 points; 7 algorithmic + 8 LLM), Quality (35 points; 35 algorithmic + 0 LLM), Feasibility (15 points; 10 algorithmic + 5 LLM), Novelty (5 points; 3 algorithmic + 2 LLM), and Diversity (10 points; 7 algorithmic + 3 LLM). The total score is the unweighted sum of all six components:

$$S_{\text{total}} = S_{\text{approach}} + S_{\text{orchestration}} + S_{\text{quality}} + S_{\text{feasibility}} + S_{\text{novelty}} + S_{\text{diversity}} \quad (\text{A1})$$

The design principle underlying this hybrid architecture is that dimensions with reliable quantitative proxies (e.g., pLDDT, ipTM, sequence identity, pairwise diversity) are evaluated algorithmically to eliminate LLM judgment variance, while dimensions requiring subjective expert-level assessment (e.g., strategic appropriateness of tool selection, adaptive reasoning quality, biological plausibility beyond sequence-level checks) are delegated to a cross-model LLM

judge panel (Section A.4). Critically, Quality — the largest single contributor (35 points) — is evaluated entirely algorithmically, ensuring that biophysical metrics are not subject to LLM hallucination or scoring bias.

Each component below is described in terms of its algorithmic sub-scores (computed deterministically) and, where applicable, its LLM judge sub-scores (computed via the panel described in Section A.4).

The Approach component (20 points) evaluates whether the agent employed a methodologically sound design pipeline. It comprises an algorithmic portion (10 points) and an LLM judge portion (10 points).

*Algorithmic sub-scores (10 points).* Function coverage (5 points) measures the fraction of expected design functions that were actually invoked: $S_{\text{coverage}} = 5 \times |F_{\text{executed}} \cap F_{\text{expected}}| / |F_{\text{expected}}|$, where $F_{\text{expected}}$ is the set of design functions specified in the task’s ground truth and $F_{\text{executed}}$ is the set of functions the agent actually called (determined by mapping each invoked MCP tool to its corresponding design functions via the **DesignFunction** enumeration). Validation inclusion (2.5 points) awards full credit if the agent included at least one validation step (structure prediction, stability scoring, or physics validation) after design generation. Iterative refinement (2.5 points) awards full credit if the agent performed at least one round of iterative refinement — defined as invoking a design or optimization function after receiving validation results. Both validation inclusion and iterative refinement are assessed by analyzing the temporal ordering of tool calls in the agent’s execution trace.

*LLM judge sub-score: Approach Strategy (10 points).* The algorithmic sub-scores capture *whether* design functions were invoked, but cannot assess *how strategically appropriate* the tool selection was for the specific task. For example, an agent that invokes **design.binder** with generic parameters and an agent that specifies epitope-specific hotspot residues for RFDiffusion conditioning both receive full function coverage credit, yet the latter reflects substantially deeper strategic reasoning. The LLM judge evaluates this dimension on a 0–10 scale using explicit score-level rubrics (Section A.4): 9–10 for target-optimised tool selection, 7–8 for appropriate selection with minor suboptimality, 5–6 for reasonable but generic strategy, 3–4 for partially appropriate with critical omissions, and 0–2 for inappropriate or random tool selection. The judge score is scaled and added to the algorithmic portion, capped at the component maximum of 20.

The Orchestration component (15 points) evaluates whether the agent executed tools in a logically correct order and performed appropriate intermediate validation. It comprises an algorithmic portion (7 points) and an LLM judge portion (8 points).

*Algorithmic sub-scores (7 points).* Pipeline ordering (4 points) checks whether design functions were invoked in a valid sequential order (e.g., backbone generation before sequence design, sequence design before structure prediction). The scoring extracts the ordered list of design functions from the tool call trace and verifies that no function was invoked before its logical prerequisites. Intermediate validation (3 points) checks whether the agent validated intermediate outputs before proceeding to subsequent steps — for example, checking structure prediction confidence (pLDDT) before proceeding to interface analysis, or verifying sequence validity before structure prediction.

*LLM judge sub-score: Orchestration Reasoning (8 points).* The algorithmic sub-scores verify structural properties of the pipeline (correct ordering, presence of validation steps) but cannot assess the quality of adaptive reasoning — whether the agent intelligently modified its strategy based on intermediate results, handled tool errors gracefully, or demonstrated scientific reasoning in its pipeline decisions. The LLM judge evaluates this dimension on a 0–8 scale: 7–8 for logical pipeline with error handling and adaptive reasoning based on intermediate results, 5–6 for correct ordering with some validation but limited adaptation, 3–4 for basic pipeline with missing validation or illogical ordering, and 0–2 for absent pipeline logic or reasoning-free tool invocation. The judge score is added to the algorithmic portion, capped at the component maximum of 15.

The Quality component (35 points; entirely algorithmic, 0 LLM) is the largest single contributor to the total score and evaluates the structural and biophysical quality of designed proteins using independently computed biophysical metrics. No LLM judgment is involved in Quality scoring, ensuring that structural assessment is free from hallucination or scoring bias. The 35 points are partitioned across three tiers — structure confidence (Tier A), interface or functional similarity (Tier B), and interface physics (Tier C) — with the partition adapting to whether the task involves a binding interface. The binding/non-binding split is determined by ground-truth thresholds (any of **ipTM\_good**, **kd\_nM\_good**, **predicted\_ddG\_good**, **active\_site\_rmsd\_good**) with a fallback to the category-specific primary metric in **QUALITY\_METRICS** (Section A.1). Concretely, binding tasks receive (Tier A, Tier B, Tier C) = (12, 18, 5) points, emphasising interface quality; non-binding tasks receive (25, 10, 0), shifting weight to monomer structure confidence and functional similarity.

All Quality sub-scores use a continuous four-band interpolation formula. Each metric is scored against three calibrated thresholds ($t_{\text{pass}}$, $t_{\text{good}}$, $t_{\text{excellent}}$), with an auto-derived floor $t_{\text{floor}} = 0.7 t_{\text{pass}}$ (for higher-is-better metrics)

that anchors the lower end of the scoring range. The fractional score $f \in [0, 1]$ is then

$$f(x) = \begin{cases} 0 & \text{if } x \leq t_{\text{floor}} \\ 0.33 \cdot \frac{x - t_{\text{floor}}}{t_{\text{pass}} - t_{\text{floor}}} & \text{if } t_{\text{floor}} < x \leq t_{\text{pass}} \\ 0.33 + 0.33 \cdot \frac{x - t_{\text{pass}}}{t_{\text{good}} - t_{\text{pass}}} & \text{if } t_{\text{pass}} < x \leq t_{\text{good}} \\ 0.66 + 0.34 \cdot \frac{x - t_{\text{good}}}{t_{\text{excellent}} - t_{\text{good}}} & \text{if } t_{\text{good}} < x \leq t_{\text{excellent}} \\ 1 & \text{if } x > t_{\text{excellent}} \end{cases} \quad (\text{A2})$$

For metrics where lower values are better (e.g., interface pAE, predicted $K_d$), the threshold ordering and floor are inverted: $t_{\text{floor}} = t_{\text{pass}}(1 + 0.3 \text{sgn}(t_{\text{pass}}))$, with the four bands traversed from $t_{\text{excellent}}$ (best) down through $t_{\text{good}}, t_{\text{pass}}, t_{\text{floor}}$. Within a tier, multiple metric sub-scores are averaged and the average is multiplied by the tier’s point allocation, giving smooth, differentiable scoring without cliff effects at threshold boundaries.

**Tier A: Structure confidence** assesses predicted monomer structural quality using two metrics: pLDDT with thresholds $(t_{\text{pass}}, t_{\text{good}}, t_{\text{excellent}}) = (65, 80, 90)$ and pTM with thresholds $(0.45, 0.65, 0.80)$. The two fractional sub-scores are averaged and multiplied by the Tier A point allocation (25 for non-binding, 12 for binding).

**Tier B: Interface or functional similarity** applies category-specific scoring. For binding tasks (Tier B = 18 points), interface quality is assessed via AlphaFold2-Multimer ipTM with thresholds $(0.15, 0.40, 0.70)$ — recalibrated to the realistic AF2-Multimer distribution observed in our task set, where the median ipTM is approximately 0.14 — and interface pAE with reversed thresholds $(25.0, 15.0, 8.0)$ (lower is better). For non-binding tasks (Tier B = 10 points), Tier B instead measures functional similarity to an oracle reference: the maximum sequence identity between any designed sequence and the published reference is mapped through the same banded scoring, awarding partial credit for designs that approach (but do not reproduce) the published solution.

**Tier C: Interface physics** (binding tasks only; 5 points) evaluates the physical plausibility of the designed binding interface. The primary signals are predicted dissociation constant $K_d$ (lower is better), interface $\Delta\Delta G$ (more negative is better), and active-site RMSD to the reference structure (lower is better), each thresholded against ground-truth values in the task definition. When ground-truth physics thresholds are unavailable, scoring falls back to interface buried surface area $(t_{\text{pass}}, t_{\text{good}}, t_{\text{excellent}}) = (800, 1500, 2500) \text{ \AA}^2$ and hydrogen-bond count $(5, 15, 30)$. Non-binding tasks receive zero Tier C points by construction.

When a task produces multiple designs, the Quality score is computed for the best-performing design (maximum across all generated sequences for each metric), reflecting the practical protein design paradigm where a researcher generates multiple candidates and selects the best for experimental validation.

The Feasibility component (15 points) assesses whether the designed protein sequences are biophysically plausible. It comprises an algorithmic portion (10 points) and an LLM judge portion (5 points).

*Algorithmic sub-scores (10 points).* Amino acid validity (4 points) checks that all designed sequences contain only the 20 standard amino acids, with partial credit for sequences containing non-standard characters. Length compliance (3 points) verifies that designed sequences fall within the length range specified in the task constraints, with graduated scoring for sequences that deviate by up to 20% from the specified range. Composition analysis (3 points) assesses whether the amino acid composition of designed sequences is consistent with naturally occurring proteins — specifically checking for anomalous over-representation of any single amino acid (e.g., $> 30\%$ of a single residue type) and ensuring reasonable hydrophobic/hydrophilic balance.

*LLM judge sub-score: Biological Feasibility (5 points).* The algorithmic sub-scores capture sequence-level validity but cannot assess higher-order biological plausibility — for example, whether CDR loop conformations are realistic for antibody tasks, whether active site geometry is consistent with catalytic function for enzyme tasks, or whether designed disulfide bond patterns are sterically feasible. The LLM judge evaluates this dimension on a 0–5 scale: 4–5 for biologically sound designs (appropriate loop geometry, consistent active site, no steric clashes), 2–3 for generally plausible with minor concerns, and 0–1 for biologically implausible designs (e.g., all-alanine core, impossible disulfide patterns). The judge score is added to the algorithmic portion, capped at the component maximum of 15.

The Novelty component (5 points) encourages agents to generate genuinely new designs rather than reproducing known sequences. It comprises an algorithmic portion (3 points) and an LLM judge portion (2 points).

*Algorithmic sub-score (3 points).* Sequence novelty is computed based on sequence identity between the designed sequence and the reference sequence provided in the ground truth. Three scoring modes are used depending on task type: for *de novo* tasks, novelty is scored inversely proportional to sequence identity to any known homolog (lower identity = higher novelty); for redesign tasks, novelty rewards designs that introduce meaningful mutations while maintaining the target fold (optimal range: 70–95% identity); and for tasks without a reference sequence, novelty is assessed based on sequence diversity relative to the input template.

*LLM judge sub-score: Novelty Quality (2 points).* Sequence identity captures surface-level novelty but cannot assess whether mutations are *meaningfully* innovative — for example, a design with 50% identity that introduces a creative new binding mode is more novel than one with 50% identity from random mutations. The LLM judge evaluates this dimension on a 0–2 scale: 2 for meaningful innovation (new fold topology, creative binding mode, non-obvious sequence strategy), 1 for some novelty but largely incremental variation, and 0 for trivially similar to the reference or random. The judge score is added to the algorithmic portion, capped at the component maximum of 5.

The Diversity component (10 points) rewards agents for generating distinct designs, reflecting the practical need for diverse candidates in experimental protein design campaigns. It comprises an algorithmic portion (7 points) and an LLM judge portion (3 points).

*Algorithmic sub-scores (7 points).* The algorithmic portion comprises two sub-components, weighted 65:35:

$$S_{\text{diversity,algo}} = 0.65 \times S_{\text{pairwise}} + 0.35 \times S_{\text{entropy}} \quad (\text{A3})$$

Pairwise sequence diversity measures the mean pairwise diversity across all generated designs:

$$S_{\text{pairwise}} = \frac{1}{\binom{N}{2}} \sum_{i < j} (1 - \text{seqid}_{ij}) \times P_{\text{max}} \quad (\text{A4})$$

where $N$ is the number of generated designs, $\text{seqid}_{ij}$ is the pairwise sequence identity between designs $i$ and $j$, and $P_{\text{max}} = 7$. *Sequence entropy* quantifies positional diversity using the mean per-position Shannon entropy normalized by $\log 20$ (the maximum entropy over 20 amino acids):

$$S_{\text{entropy}} = \frac{1}{L} \sum_{k=1}^L \frac{H_k}{\log 20} \times P_{\text{max}}, \quad H_k = - \sum_{a=1}^{20} p_{a,k} \log p_{a,k} \quad (\text{A5})$$

where $L$ is the aligned sequence length, $H_k$ is the Shannon entropy at position $k$, and $p_{a,k}$ is the frequency of amino acid $a$ at position $k$. We deliberately exclude a design count sub-component: because task prompts do not specify how many designs to generate, rewarding count would unfairly penalize agents (e.g., human experts) that produce fewer but higher-quality candidates.

*LLM judge sub-score: Diversity Quality (3 points).* Pairwise sequence identity and Shannon entropy capture statistical diversity but cannot assess whether the diversity is *functionally meaningful* — for example, whether designs explore different binding modes, conformational strategies, or topological folds, as opposed to superficial sequence variation that preserves the same structural solution. The LLM judge evaluates this dimension on a 0–3 scale: 3 for functionally diverse designs exploring distinct binding modes, conformations, or strategies; 1–2 for some diversity but largely minor variants of a single approach; and 0 for no meaningful diversity (single design or near-identical copies). The judge score is added to the algorithmic portion, capped at the component maximum of 10.

### A.4 LLM Judge Panel

The 28 LLM-assessed points across five rubric dimensions (Approach Strategy 10, Orchestration Reasoning 8, Biological Feasibility 5, Novelty Quality 2, Diversity Quality 3) are evaluated by a cross-model panel of LLM judges following the Panel of LLM Evaluators (PoLL) principle [2]. This section describes the judge panel composition, prompt structure, score aggregation, and reliability mechanisms.

#### A.4.1 Judge panel composition and self-exclusion

For each evaluated agent, three judges are drawn from four candidate model families: Claude Sonnet 4 (Anthropic), GPT-5.2 (OpenAI), Gemini 2.5 Pro (Google), and DeepSeek (DeepSeek). Critically, the evaluated agent’s own model family is excluded from its judge panel to mitigate self-preference bias — the tendency of LLMs to systematically favour their own outputs over those of other models [3]. For example, outputs from Claude Sonnet 4.5 agents are judged by GPT-5.2, Gemini 2.5 Pro, and DeepSeek; outputs from GPT-5 agents are judged by Claude Sonnet 4, Gemini 2.5 Pro, and DeepSeek. Human baselines (Expert and Oracle) are evaluated by all four judges, as no self-preference bias applies.

#### A.4.2 Judge prompt structure

Each judge receives a structured prompt comprising six sections: (1) *Task Description* — the original design objective, target protein specification, and success criteria; (2) *Reference Pipeline* — the expert-verified optimal tool sequence for the task (e.g., RFdiffusion → ProteinMPNN → AlphaFold2), providing context for evaluating the agent’s strategic choices; (3) *Agent’s Tool Call Log* — the numbered sequence of tool invocations with arguments, representing the agent’s execution trace; (4) *Designed Sequences* — up to 10 designed sequences in FASTA format (truncated to 80 characters per line); (5) *Algorithmic Metrics* — pLDDT, ipTM, predicted $K_d$, and other quantitative metrics provided as read-only context; and (6) *Scoring Rubric* — the full text of all five LLM-assessed dimensions with explicit score-level descriptions, plus a JSON output format specification.

Algorithmic metrics are provided to the judge as read-only context with explicit instructions not to use them for scoring: “The following metrics are provided for biological context only. Do NOT use these numbers to adjust your scores on any dimension. Your evaluation should be based solely on the agent’s reasoning, tool selection, and design strategy as described in the rubric.” This design prevents the halo effect — where, for example, seeing ipTM = 0.45 might cause a judge to lower Approach Strategy scores despite the agent having selected an appropriate pipeline.

#### A.4.3 Score aggregation

Judge scores are aggregated via outlier-robust weighted averaging. For each of the five LLM-assessed dimensions independently:

1. *Median computation*: the median of the three judge scores is computed.
2. *Outlier detection*: any judge score deviating more than 2 points from the median is flagged as an outlier.
3. *Weighted average*: normal scores receive weight $w = 1.0$; outlier scores receive weight $w = 0.5$ (downweighted but not discarded, preserving information while reducing the influence of aberrant scores).
4. *Range clamping*: the weighted average is clamped to $[0, s_{\max}]$, where $s_{\max}$ is the dimension’s maximum score.

For example, if three judges assign Approach Strategy scores of $[8, 7, 3]$: median = 7; the score of 3 deviates by $|7 - 3| = 4 > 2$, so it is flagged as an outlier; weighted average = $(8 \times 1.0 + 7 \times 1.0 + 3 \times 0.5) / (1.0 + 1.0 + 0.5) = 16.5 / 2.5 = 6.6$.

#### A.4.4 Hybrid score merging

The algorithmic and LLM judge scores are merged per component as follows. First, the algorithmic score is computed from tool call logs, sequences, and biophysical metrics as described in the component definitions above. Second, the aggregated LLM judge score for each applicable dimension is added to the algorithmic score. Third, the merged score is capped at the component’s rubric maximum to prevent double-counting (e.g., if the algorithmic Approach score is 8.0 and the judge Approach Strategy score is 8.5, the merged Approach score is $\min(8.0 + 8.5, 20) = 16.5$). For Quality, no merging is needed as the LLM portion is zero.

#### A.4.5 Reliability mechanisms

Several mechanisms ensure scoring reliability beyond self-exclusion and outlier downweighting:

• *Response parsing robustness*: judge outputs are parsed using a three-stage fallback: JSON embedded in markdown code blocks → direct JSON parsing → regex extraction of score fields. If all three fail, the dimension receives the midpoint score (half of maximum), ensuring that API failures do not produce zero scores.
• *Score clamping*: any judge score outside the valid range $[0, s_{\max}]$ is replaced with the midpoint score for that dimension.
• *Rubric cap enforcement*: the merged algorithmic + LLM score for each component is capped at the component maximum, preventing the LLM judge from effectively “double-scoring” aspects already captured by the algorithmic portion.
• *Continuous scoring with linear interpolation*: within each score-level band (e.g., 5–6, 7–8), judges are instructed to assign continuous scores rather than integers, reducing cliff effects at band boundaries.
• *Dry-run mode*: for testing and validation, a deterministic dry-run mode returns midpoint scores for all dimensions without making API calls, enabling end-to-end pipeline testing.

### A.5 Independent Quality Verification

To ensure that Quality scores (Section A.3) reflect actual structural and biophysical properties rather than agent-reported metrics, all designed sequences undergo independent verification using AlphaFold2 [4] post-evaluation. This is critical because agents may report inflated or incorrect quality metrics: an agent might claim high pLDDT based on a different prediction model, misparse tool outputs, or hallucinate confidence values. By re-evaluating all designs with a single, standardized AlphaFold2 pipeline, we obtain unbiased quality assessments that are comparable across all agents and conditions.

The post-evaluation pipeline operates as follows. For each agent’s output on each task, we extract all designed protein sequences (filtering out invalid sequences containing non-amino-acid characters). For non-binding tasks, each sequence is submitted to AlphaFold2 monomer prediction (ColabFold v1.5.5, 3 recycles, no templates) to obtain pLDDT and pTM scores. For binding tasks, each designed sequence is combined with the target chain(s) specified in the task definition and submitted to AlphaFold2-Multimer prediction to obtain ipTM, interface pAE, and per-chain pLDDT scores; additionally, the predicted complex structure undergoes interface analysis using PyRosetta to compute binding $\Delta\Delta G$ (via the `InterfaceAnalyzerMover`), buried surface area (BSA), and interface contact maps. All post-evaluation computations are performed on NVIDIA A100 GPUs with identical random seeds to ensure reproducibility.

Post-evaluation coverage across conditions is near-complete. In unguided mode: GPT-5 64/64 tasks (100%), Sonnet 4.5 60/60 tasks (100%), DeepSeek V3 73/73 tasks (100%), and Gemini 2.5 Pro 59/59 tasks (100%). In guided mode: GPT-5 66/68 tasks (97%), Sonnet 4.5 61/63 tasks (97%), DeepSeek V3 73/74 tasks (99%), and Gemini 2.5 Pro 61/61 tasks (100%). The hardcoded pipeline achieved 48/48 tasks (100%). Tasks without post-evaluation results (due to sequence extraction failures or timeout) receive zero Quality points for the affected sub-components.

### A.6 Data Contamination Defense

Because all benchmark tasks are derived from published protein design studies (2024–2025), there is a risk that LLM agents could retrieve published solutions from their training data or through web search rather than performing genuine design. We implement a five-layer defense-in-depth strategy to prevent and detect such contamination.

Before any task is presented to an agent, the task sanitizer (`sanitizer.py`) strips six metadata fields that could enable contamination: `source` (the paper title and authors), `doi` (the Digital Object Identifier), `tools_expected` (the expected tool pipeline), `tags` (categorical labels), `test_file` (path to validation scripts), and `ground_truth` (reference sequences and structures). The sanitized task retains only the design specification (description, constraints, input PDB, target/design chains) — sufficient for an agent to attempt the design but insufficient to identify the source publication.

Every agent system prompt includes an explicit anti-contamination instruction: “You are a protein design assistant. Design proteins using ONLY the provided MCP tools. NEVER search for, reference, or use information from published papers, preprints, DOIs, or known protein design solutions. All designs must be generated *de novo* using the computational tools available to you. If you recognize a task from published literature, you must still solve it independently using only the provided tools.” This instruction is appended to all agent configurations regardless of the underlying LLM.

All agent executions occur within Docker containers with network access disabled (`--network=none`). This prevents agents from making HTTP requests to search engines, academic databases (PubMed, bioRxiv), or any external service during task execution. The only I/O channels available to the agent are the MCP tool interface (via stdio) and the local filesystem within the container.

Ground truth files (reference sequences, structures, and expected metrics stored in `data/oracle/sequences.json` and per-task `ground_truth/` directories) are mounted in a separate Docker volume that is not accessible from the agent’s execution environment. The scoring pipeline accesses ground truth only during post-hoc evaluation, never during agent execution.

After all runs are complete, a contamination detection module (`contamination.py`) scans each agent’s full reasoning trace (all text generated by the LLM during task execution) for four contamination signals: (1)DOI mentions — any string matching the DOI pattern `10.\d{4,}/` triggers a positive signal; (2)Source paper references — mentions of specific author names, paper titles, or journal names associated with the task’s source publication; (3)Known sequence reproduction — designed sequences with $\geq 95\%$ identity to the ground truth reference sequence (for tasks where random chance of such similarity is negligible); and (4)Methodology leakage — specific methodological details from the source paper (e.g., exact parameter values, experimental conditions) that appear in the agent’s reasoning but were not provided in the task prompt. Each signal contributes to a composite contamination score $c \in [0, 1]$. If $c \geq 0.5$ for any task, the agent’s score for that task is set to zero (cheating penalty). We report the number of flagged tasks per agent in the supplementary results.

### A.7 Experimental Setup

We evaluate four frontier LLMs as protein design agents. **GPT-5** (OpenAI; model string `gpt-5.2`) is accessed via the OpenAI Python SDK using its native function-calling interface. **Claude Sonnet 4.5** (Anthropic; model string `claude-sonnet-4-5-20250929`) is accessed via the Anthropic Python SDK. **DeepSeek V3** (DeepSeek; model string `deepseek-chat`) is accessed via the OpenAI-compatible endpoint at https://api.deepseek.com. **Gemini 2.5 Pro** (Google; model string `gemini-2.5-pro`) is accessed via Vertex AI through the Google Gen AI Python SDK with the explicit `thinking_budget=0` setting so its reasoning mode is disabled, ensuring a fair comparison against the other

non-thinking baselines. MCP tool definitions are translated to each SDK’s native tool schema format on the fly by the BioDesignBench tool provider (one of `get_tool_definitions_anthropic` / `openai` / `gemini`). Each agent session is allowed up to 50 tool-use turns (`max_iterations=50`); individual tool invocations run inside a Docker container with a 300-second wall-clock timeout per call.

All agents share a common architecture: the LLM receives a system prompt (including the anti-contamination instruction and mode-specific guidance from `_BENCHMARK_GUIDANCE` or `_USER_GUIDANCE`), the task prompt, and the active MCP tool catalog (16 atomic tools in unguided/benchmark mode or all 19 tools in guided/user mode). The agent then engages in a multi-turn conversation with the MCP server, alternating between reasoning and tool invocation; tool results are returned as structured JSON and appended to the conversation history. The session terminates when the agent produces a final output (designed sequences plus a `metrics.json` file), reaches `max_iterations`, or hits an SDK-level error. Each agent is registered with three mode-aware identifiers in `biodesignbench/agents/_init_.py`: a base ID inheriting the run-level `--tool-mode` flag (e.g., `gpt5-tools`), a benchmark-mode variant (e.g., `gpt5-tools-benchmark`, equivalent to “unguided” in the paper), and a user-mode variant (e.g., `gpt5-tools-user`, equivalent to “guided”).

The hardcoded pipeline baseline (`biodesignbench/agents/baselines/hardcoded_pipeline.py`) executes a deterministic, LLM-free tool chain for each task by dispatching on the task’s (`DesignApproach`, `MolecularSubject`) category from `taxonomy.get_category()`, with a special case that routes any task whose ID begins with `cfid_` into the conformational/functional design pipeline regardless of subject. Five pipeline variants are defined: (1) *De novo binder* (`_pipeline_dnb`, used for de novo antibody/binder tasks): `suggest_hotspots` (only when no hotspot residues are provided) → `design_binder` → `validate_design`; (2) *Sequence optimisation* (`_pipeline_sqo`, used for all redesign tasks): structure prefetch via `predict_structure` (only when no input PDB exists) → `design_sequence` (ProteinMPNN, temperature 0.2) → `score_stability`; (3) *De novo unconditional* (`_pipeline_dnk`, used for unconditional scaffold tasks without binding-site residues): `generate_backbone` → `design_sequence` (ProteinMPNN, temperature 0.1, $n = 4$, best-by-pLDDT selection); (4) *Complex/interface engineering* (`_pipeline_cpx`, used for de novo enzyme/scaffold tasks with a target interface): `suggest_hotspots` (when needed) → `design_binder` (with `generate_backbone` → `design_sequence` as a fallback) → `predict_complex`; (5) *Conformational/functional design* (`_pipeline_cfd`, used for de novo enzyme/fluorescent-protein tasks and any `cfid.*` task): `predict_structure` → `design_sequence` → `energy_minimize` → `validate_design`. Each variant uses default parameters without task-specific tuning, providing a deterministic reference for the performance achievable by a non-adaptive system.

A human expert (first author, with 3+ years of computational protein design experience) independently solves each task using the same 17 MCP tools available to the LLM agents. The expert has full access to tool documentation and can freely choose which tools to invoke, in what order, and with what parameters. The expert also has access to domain knowledge about protein design best practices but is prohibited from consulting the source publications for any task. Five representative pipeline strategies are employed across tasks: (1) RFdiffusion pipeline: `design_binder` or `generate_backbone` → `optimize_sequence` (8 samples, temperature 0.2) → `predict_structure` → `analyze_interface`; (2) ProteinMPNN optimization: `optimize_sequence` (16 samples, temperature 0.1) → `predict_structure` → `score_stability` with iterative refinement; (3) Rosetta design: `rosetta_relax` → `rosetta_design` → `predict_structure` → `rosetta_interface_score`; (4) ESM-guided evolution: `optimize_sequence` → `predict_structure` → `score_stability` (iterative, 3 rounds); and (5) Hybrid pipeline: combining multiple tools with task-specific parameter tuning. The expert baseline provides an upper bound reference for what is achievable with domain expertise and the same tool set.

The human oracle baseline represents the theoretical ceiling for each task. For each task, the ground truth reference sequence (obtained from the source publication) is submitted to AlphaFold2 post-evaluation to obtain quality metrics. The oracle’s Approach, Orchestration, Feasibility, and Diversity scores are set to maximum values, and the Quality score is computed from the AF2 metrics of the published sequence. The Novelty score is set to zero (as the oracle sequence is by definition the reference). Oracle scoring is computed by measuring sequence identity between the published sequence and itself (yielding identity = 1.0) and applying the standard scoring rubric with oracle-specific overrides. This baseline quantifies the maximum achievable Quality score for each task given the published solution.

All benchmark runs are executed on a compute cluster with 8× NVIDIA A100 (80GB) GPUs. Agent sessions are parallelized using Python’s `asyncio.gather` with one GPU assigned per concurrent session (`--gpu-id` flag). The benchmark runner (`run_benchmark.py`) accepts command-line arguments for agent selection (`--agent`), task tier (`--tier tier2`), MCP server path (`--mcp-server`), Docker execution (`--docker`), GPU assignment (`--gpu-id`), per-task timeout (`--timeout 120`), tool mode (`--tool-mode unguided|guided`), and run resumption (`--resume <run_id>`). The resume functionality enables recovery from interrupted runs by checkpointing completed tasks and skipping them on restart. A complete benchmark evaluation (4 LLMs × 2 modes × 76 tasks = 608 agent sessions, plus 76 hardcoded pipeline runs) requires approximately 72 GPU-hours.

### Appendix B Statistical Analysis

To assess whether performance differences among the 11 evaluation conditions are statistically robust, we conduct non-parametric hypothesis testing on per-task total scores. Non-parametric tests are chosen because score distributions across conditions violate the assumptions of parametric alternatives: the Oracle distribution is left-skewed near the scoring ceiling, Gemini 2.5 Pro distributions are right-skewed near the floor, and a Levene test rejects homoscedasticity ($p < 0.001$). All statistical analyses are performed in Python using SciPy v1.11 [5] and pingouin v0.5 [6].

#### B.1 Omnibus test

A Kruskal–Wallis $H$ test [7] across all 11 conditions ($n = 76$ tasks per condition; 836 total observations) yields $H = 384.6$ ($df = 10$, $p < 10^{-76}$), decisively rejecting the null hypothesis that all conditions share the same score distribution. We therefore proceed with pairwise comparisons to identify which specific condition pairs differ.

#### B.2 Pairwise comparisons

For between-condition comparisons, we use two-sided Mann–Whitney $U$ tests [8], which compare rank sums without assuming normality. All $p$-values are corrected for multiple comparisons using the Bonferroni method [9] with $m = 55$ pairwise tests, yielding a corrected significance threshold of $\alpha = 0.05/55 \approx 9.1 \times 10^{-4}$. Although the Bonferroni correction is conservative (increasing the risk of Type II errors), we prefer it over less stringent alternatives (e.g., Benjamini–Hochberg) because we report the full $11 \times 11$ matrix and wish to minimise false-positive claims of significance.

Supplementary Figure S1 presents the full pairwise statistical comparison of all 11 conditions. Panel (a) displays a $-\log_{10}(p)$ heatmap of two-sided Mann–Whitney $U$ tests with Bonferroni correction for 55 comparisons. Panel (b) shows Cohen’s $d$ effect sizes for each pair. Three tiers of separation emerge: (i) Oracle is significantly different from all other conditions ($p < 0.001$ for all 10 pairs; Oracle mean = 74.9); (ii) Human Expert (61.3) and DeepSeek V3 unguided (60.4) occupy a second tier with broadly overlapping confidence intervals, joined by DeepSeek V3 guided (58.5), GPT-5 unguided (55.6), GPT-5 guided (55.3), and the Hardcoded Pipeline (54.2); (iii) Sonnet 4.5 shows a wider mode gap (guided 50.2, unguided 41.2), while Gemini 2.5 Pro defines the lower bound (guided 8.8, unguided 8.1). The two Gemini modes are indistinguishable ($p = 0.972$), confirming that enhanced tool descriptions provide no benefit when the underlying tool-calling mechanism fails entirely.

#### B.3 Effect sizes

To complement $p$-values with measures of practical significance, we compute rank-biserial correlations ($r_{rb}$) [10] for the within-model guided-vs.-unguided mode comparisons. Sonnet 4.5 is the only model with a statistically significant guided-mode advantage ($r_{rb} = -0.325$, $p = 5.40 \times 10^{-4}$), representing a medium effect. GPT-5 ($r_{rb} = -0.073$, ns), DeepSeek V3 ($r_{rb} = -0.029$, ns), and Gemini 2.5 Pro ($r_{rb} = +0.064$, ns) all show negligible effects on total score. The selective significance of Sonnet 4.5 suggests that guided-mode hints are most beneficial for models at an intermediate competency level — strong enough to act on richer descriptions, but not so strong as to already saturate the available tool-use strategies.

### Appendix C Scoring Rubric Validation

A critical concern for any multi-component scoring rubric is whether condition rankings are robust to reasonable perturbations in component weights. If small weight changes produced substantially different rankings, the benchmark would be sensitive to arbitrary design choices rather than reflecting genuine performance differences.

Supplementary Figure S2 displays a weight perturbation analysis in which each of the six quality sub-metrics used in the Quality scoring component (pLDDT, pTM, Ramachandran, Rosetta, ESM-2 PPL, PAE) is individually scaled by multipliers ranging from $0.5\times$ to $2.0\times$ while all other weights remain at their default values. For each perturbation we recompute condition-level total scores and measure the Spearman $\rho$ between the perturbed and default condition rankings. All perturbations yield $\rho \geq 0.973$, with the vast majority achieving $\rho = 1.000$ (perfect rank preservation). This extreme stability indicates that no single sub-metric dominates the Quality score to the extent that its reweighting could alter condition rankings, providing strong empirical support for the robustness of the rubric.

We additionally verified that the Approach and Orchestration components contribute orthogonal information to Quality. Recomputing Approach and Orchestration scores for 50 randomly sampled outputs yielded perfect agreement with the stored values (Pearson $r = 1.000$ for both components), confirming reproducibility of the rubric implementation.

### Pairwise Statistical Comparisons

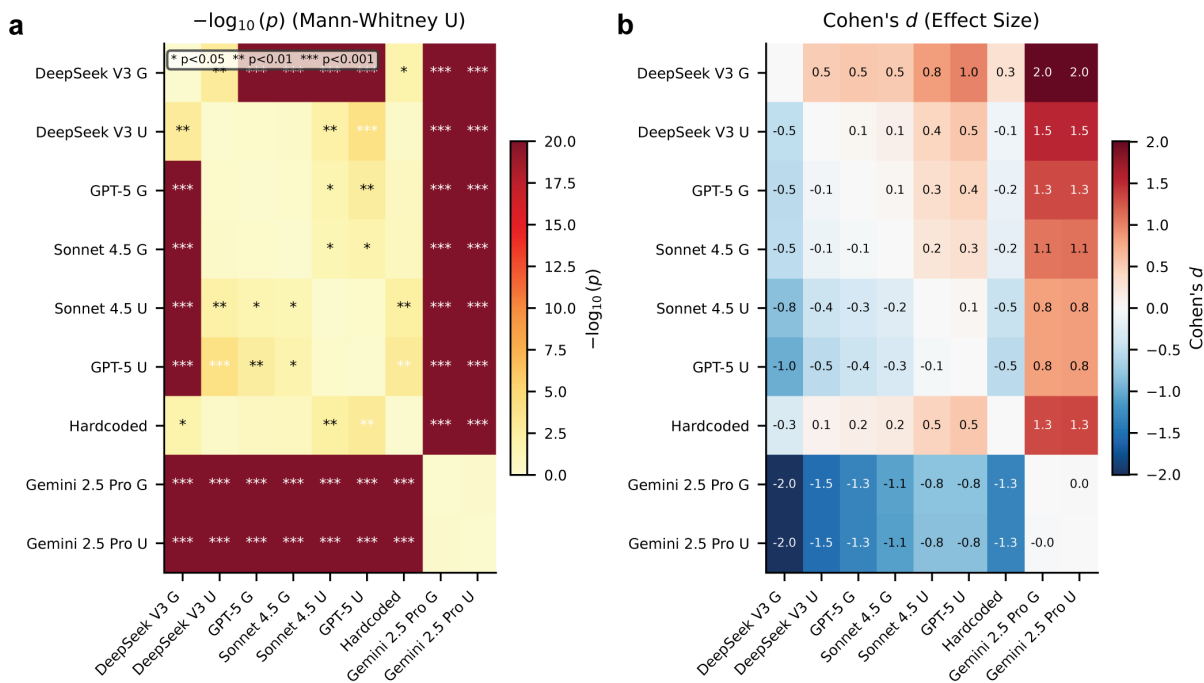

**Fig. S1 Pairwise statistical comparison of all 11 conditions.** (a) $-\log_{10}(p)$ heatmap of two-sided Mann-Whitney $U$ tests between all condition pairs, with Bonferroni correction for 55 comparisons ($\alpha = 9.1 \times 10^{-4}$). Annotations: \*$p < 0.05$, \*\*$p < 0.01$, \*\*\*$p < 0.001$; ns indicates not significant after correction. (b) Cohen's $d$ effect size matrix. Positive values (red) indicate that the row condition outperforms the column condition. The Oracle tier ($d > 1.0$ vs. all others), the mid-tier convergence among Expert/DeepSeek/GPT-5/Hardcoded, and the Gemini floor ($d > 1.5$ vs. all non-Gemini conditions) are clearly visible.

### Appendix D Independent Structural Verification

Quality scores rely on metrics obtained through independent post-evaluation of all designed sequences using Boltz-2 [11], ensuring that reported quality reflects actual structural and biophysical properties rather than agent-reported (potentially hallucinated) metrics.

Supplementary Figure S3 presents multi-metric quality convergence across conditions. Panels (a)–(c) show distributions of three independent structural metrics — Boltz pLDDT, ESM-2 perplexity (PPL), and Rosetta energy per residue — across all conditions. All three metrics show consistent stratification: conditions that score well on one metric tend to score well on the others, confirming that the Quality rubric captures genuine structural quality rather than artifacts of any single evaluation tool. Panels (d)–(f) display pairwise correlations between these metrics (pLDDT vs. ESM-2 PPL: $\rho = -0.39$; pLDDT vs. Rosetta: $\rho = -0.67$; ESM-2 PPL vs. Rosetta: $\rho = 0.42$), confirming moderate cross-metric agreement with the expected sign conventions.

Median Boltz pLDDT values are closely matched among the top conditions (Oracle = 0.928, Human Expert = 0.935, DeepSeek V3 guided = 0.932; Mann-Whitney $U$: Oracle vs. Expert $p = 0.332$, ns; Oracle vs. DeepSeek $p = 0.515$, ns), indicating that the top-performing LLM agents produce designs with structural confidence comparable to both the Expert and the Oracle's published sequences. Because per-design structural quality metrics do not differentiate these conditions, the Expert's advantage in total score arises from Approach, Orchestration, and iterative refinement rather than per-design structural quality — a finding that motivates the depth analyses in subsequent sections.

### Appendix E Data Contamination Defense

To ensure that LLM agents do not achieve high scores by retrieving published solutions rather than orchestrating computational tools, we implement the multi-layered contamination defense described in Section A.6 and verify its effectiveness post hoc through a five-layer audit.

Supplementary Figure S4 displays the contamination audit results across all 76 task outputs for each of the 8 LLM conditions. The five audit layers are: (i) DOI/URL pattern matching against the source-publication metadata

### Ranking Stability Under Weight Perturbation

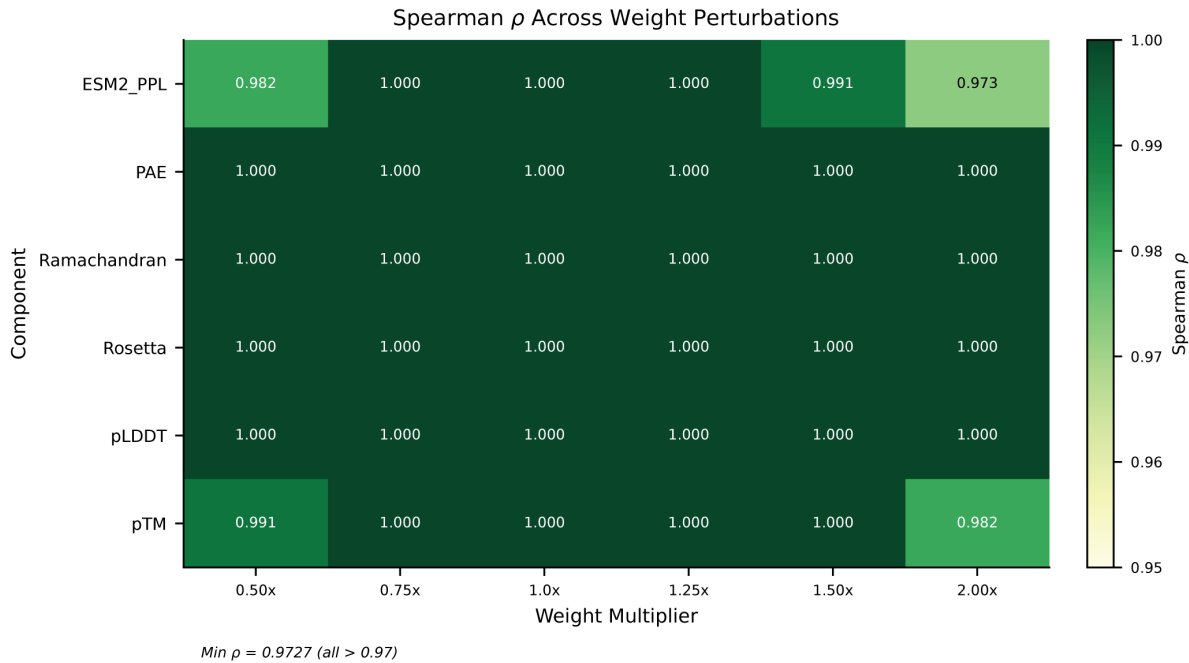

**Fig. S2 Ranking stability under scoring weight perturbation.** Heatmap of Spearman $\rho$ between the default condition ranking and rankings obtained under systematic perturbation of individual quality sub-metric weights. Rows correspond to the six sub-metrics (ESM-2 PPL, PAE, Ramachandran, Rosetta, pLDDT, pTM); columns correspond to weight multipliers (0.5× to 2.0×). All $\rho \geq 0.973$, confirming that condition rankings are robust to at least 2× changes in any individual weight.

stripped by the task sanitizer; (ii) author-name and journal-title string matching; (iii) $\geq 95\%$ sequence identity between any designed sequence and the ground-truth reference sequence; (iv) $n$-gram overlap detection between the agent’s reasoning trace and the source paper’s abstract or methods section; and (v) explicit pattern matching for known solution shortcuts (e.g., copy-pasted PDB identifiers from the source publication). Zero tasks were flagged across all conditions (0/76 per agent in every layer), confirming that the anti-contamination prompt instructions, combined with network isolation within the Docker execution environment, effectively prevented data leakage.

### Appendix F Task Stratification

To assess whether benchmark performance varies systematically by task characteristics, we stratify results along three axes: design approach (*de novo* vs. redesign), assigned difficulty level (easy, medium, hard), and molecular subject (antibody, enzyme, binder, scaffold, fluorescent protein).

#### F.1 Difficulty scaling

Supplementary Figure S5 displays mean total scores as a function of assigned difficulty level (easy: $n = 21$; medium: $n = 28$; hard: $n = 27$). Top-tier LLM agents (DeepSeek V3, GPT-5) maintain relatively stable performance across difficulty levels, with less than 5-point drops from easy to hard tasks. Sonnet 4.5 shows a steeper decline, particularly in unguided mode, suggesting greater sensitivity to task complexity. Gemini 2.5 Pro scores remain near the floor across all difficulty levels, consistent with its systematic tool-calling failure rather than task-specific limitations. The Hardcoded Pipeline shows a slight improvement on hard tasks, likely reflecting the higher proportion of structured binder design tasks in the hard category.

#### F.2 De novo versus redesign

Supplementary Figure S6 displays performance stratification by design approach. Panel (a) shows mean total scores for *de novo* ($n = 47$) and redesign ($n = 29$) tasks across all 11 conditions. *De novo* tasks yield slightly higher mean scores for most conditions, reflecting the strong alignment between the core RFdiffusion–ProteinMPNN pipeline and

### Multi-Metric Quality Convergence

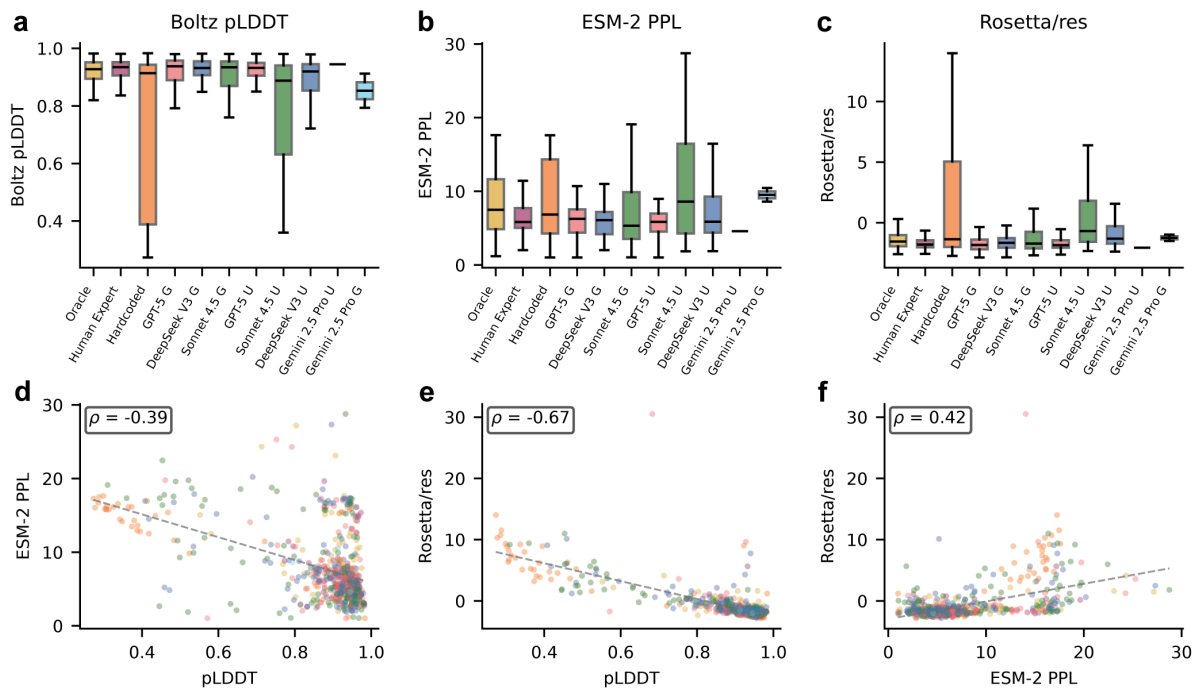

**Fig. S3 Multi-metric quality convergence across independent structural verifiers.** (a–c) Distributions of three independent structural metrics across all conditions: (a) Boltz pLDDT, (b) ESM-2 perplexity, (c) Rosetta energy per residue. Consistent stratification confirms that the Quality rubric captures genuine structural quality. (d–f) Pairwise scatter plots between structural metrics with Spearman $\rho$ annotations: (d) pLDDT vs ESM-2 PPL ($\rho = -0.39$), (e) pLDDT vs Rosetta ($\rho = -0.67$), (f) ESM-2 PPL vs Rosetta ($\rho = +0.42$), demonstrating moderate cross-metric agreement with expected sign conventions.

*de novo* design workflows. Panel (b) quantifies the approach gap ($\Delta = \text{de novo} - \text{redesign}$) per condition with Mann–Whitney significance annotations, showing that the gap is statistically significant for Oracle, Human Expert, and select LLM conditions but not for most agents, confirming that approach-level differences are modest relative to between-condition variation.

#### F.3 Molecular subject stratification

Supplementary Figure S7 displays per-task total score distributions broken out by molecular subject (antibody, enzyme, binder, scaffold, fluorescent protein) for all 11 conditions. Three qualitative patterns emerge. First, binder tasks ($n = 19$) yield the highest median scores across nearly all conditions, reflecting strong alignment between the core RFdiffusion–ProteinMPNN–AlphaFold2 pipeline and *de novo* binder design. Second, antibody tasks ($n = 9$) and fluorescent-protein tasks ($n = 11$) are the most challenging across conditions, consistent with antibody design requiring specialised CDR-loop handling and fluorescent-protein design depending on chromophore–environment interactions that current tools model only indirectly. Third, the Expert’s advantage over LLM agents is largest on antibody and fluorescent-protein tasks — precisely the domains where evaluation depth matters most — providing a qualitative preview of the depth-bottleneck analysis presented in Sections H–K. The full 11-condition boxplot with all five molecular subjects is shown in Supplementary Figure S7; this panel complements the approach- and difficulty-stratifications above by showing that domain identity, not just task properties, modulates the agent–expert gap.

### Appendix G Failure Mode Taxonomy under Guided Mode

Beyond total score, each task–condition outcome can be classified into a five-level behavioural failure-mode taxonomy that separates competence, capability, and chance. We define: **deep success** — the agent selects the correct tool pipeline, executes it successfully, and reaches an execution depth at or above the expert median; **shallow success** — correct plan and successful execution, but evaluation depth below the expert median; **tool gap** — correct plan, but execution fails (API error, parameter misuse, or tool invocation exception); **science gap** — incorrect plan (wrong tool category, missing required stage, or scientifically implausible ordering); **serendipity** — a scientifically incorrect plan

### Data Contamination Check

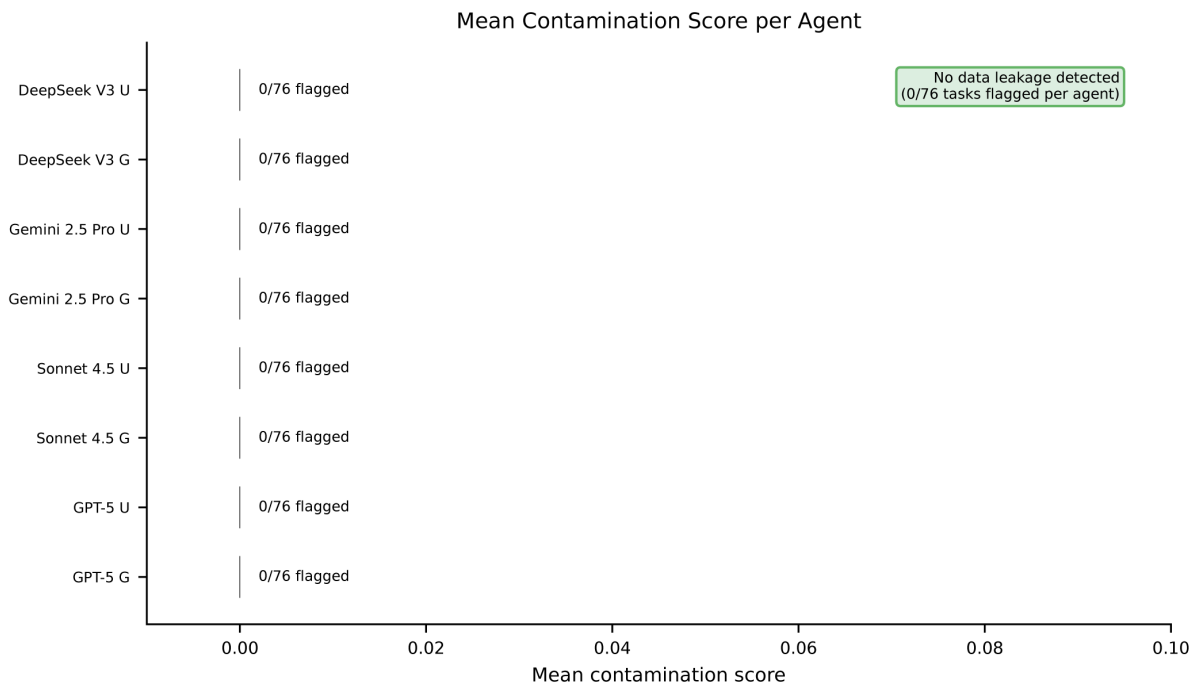

**Fig. S4 Data contamination audit.** Mean contamination score per agent across all 76 tasks for each of the 8 LLM conditions, aggregating five detection layers (DOI matching, author/title matching, sequence identity $\geq 95\%$, $n$-gram overlap with source paper, known-solution shortcuts). All conditions show 0/76 flagged tasks, confirming that the combination of metadata sanitization, anti-contamination prompt instructions, and network-isolated Docker execution effectively prevented data leakage during agent evaluation.

that nonetheless produces a high-scoring output through an emergent shortcut. The deep/shallow split is re-used in the depth analysis (main text Section 2.4; Supplementary Section H) to subdivide successful runs into depth-stratified bands.

Supplementary Figure S8 displays the per-task failure-mode decomposition for the four LLM agents under guided mode across all 76 tasks. The four stacked bars reveal distinct behavioural signatures. DeepSeek V3 achieves the highest deep-success rate (14.5% of tasks) and is otherwise dominated by shallow success and tool gap — indicating that when it fails, it fails at execution rather than at planning. GPT-5 shows a similar dominance of these two categories, with a lower deep-success fraction. Sonnet 4.5 shifts its failure mass from tool gap into science gap, consistent with its benefit from guided-mode workflows that narrow the plan search space. Gemini 2.5 Pro is dominated almost entirely by science gap, reflecting its systematic failure to invoke protein-design tools through the MCP interface (Section B); the small serendipity component corresponds to trivially lucky outputs on the few tasks where no tool use is strictly required. Across all four models, serendipity contributes negligibly ($< 5\%$ of tasks in every condition), confirming that the benchmark does not reward scientifically incorrect strategies that happen to achieve high scores.

### Appendix H Successful Runs Decomposed by Execution Depth

Supplementary Figure S9 extends the failure-mode taxonomy view (Section G) by decomposing successful runs into deep and shallow execution across all 11 conditions including the human expert and hardcoded baselines. The human expert achieves deep success on 53.8% of tasks; the strongest LLM (DeepSeek V3 unguided) reaches deep success on only 14.5% — a $3.7\times$ gap that is invisible in binary success/failure analysis. The hardcoded pipeline shows a moderate deep-success rate from its built-in evaluation loop. Gemini conditions are dominated by tool gap, consistent with their MCP tool-calling failure.

### Appendix I Volume Axis of Evaluation Effort

Main-text Fig. 3d uses the *variety* axis (number of distinct evaluation metric categories per candidate) to relate evaluation behaviour to total score. Here we report the parallel correlation along the *volume* axis (number of evaluation

### Score Scaling with Task Difficulty

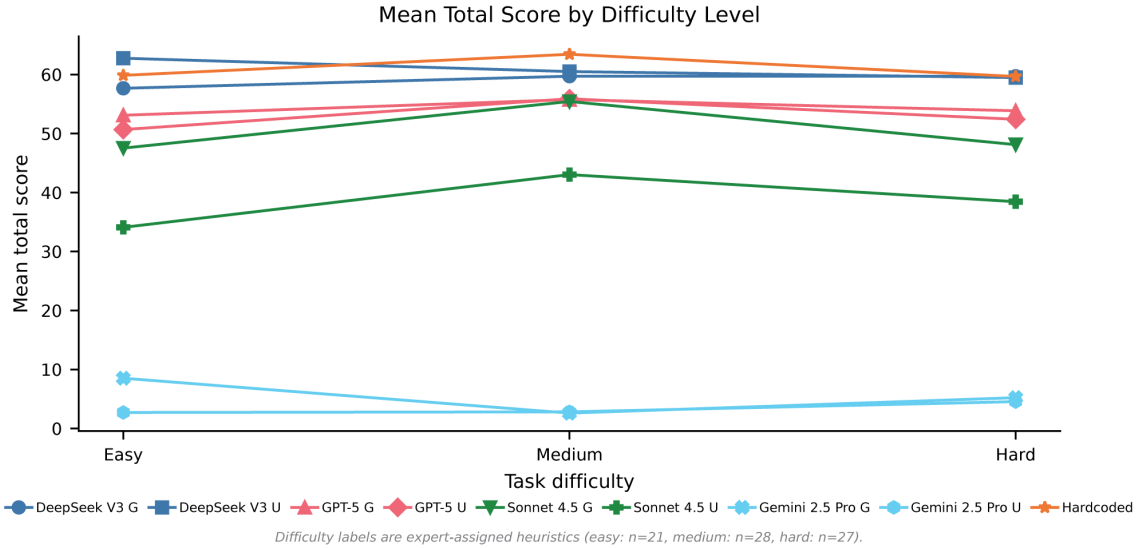

**Fig. S5 Score scaling with task difficulty.** Mean total score as a function of expert-assigned difficulty level (easy: $n = 21$; medium: $n = 28$; hard: $n = 27$) for all 11 conditions. Top-tier agents (DeepSeek V3, GPT-5) maintain stable performance across difficulty levels. Sonnet 4.5 shows steeper decline with increasing difficulty. Gemini 2.5 Pro remains near the floor at all levels.

tool calls per candidate). Plotting total score against $\log(1 + \text{evaluation calls per candidate})$ across all 836 task-condition observations yields Spearman $\rho = 0.69$, $p < 10^{-116}$ — a correlation strength similar to the variety-axis result (Supplementary Fig. S10). Volume and variety are mathematically related but distinct quantities: an agent can call AlphaFold six times (volume = 6, variety = 1) or call AlphaFold, Rosetta, and DSSP once each (volume = 3, variety = 3). The variance-partitioning analysis below shows that the variety axis carries more unique explanatory power once binary tool coverage is accounted for.

### Appendix J Stage Coverage Analysis

To examine how agents distribute their tool calls across the canonical protein-design pipeline (backbone generation → sequence design → structure prediction → scoring/validation → refinement), we analysed per-stage invocation patterns across all 836 task-condition observations.

Supplementary Figure S11 displays a stage-depth delta heatmap showing the difference in per-stage depth between guided and unguided modes ($\Delta\text{depth} = \text{guided} - \text{unguided}$) for each LLM family across four canonical pipeline stages. Two patterns emerge. First, Sonnet 4.5 shows the largest guided-mode depth gains, particularly in structure prediction (+1.28) and backbone generation (+0.19), consistent with its statistically significant guided-mode advantage in total score. Second, GPT-5 shows a notable guided-mode *decrease* in structure prediction (−0.90), suggesting that enhanced tool descriptions may redirect GPT-5’s effort away from structure prediction towards other stages. DeepSeek V3 and Gemini 2.5 Pro show minimal mode effects across all stages.

Supplementary Figure S12 decomposes the total score change under guidance into individual rubric components for each LLM family. Sonnet 4.5’s large total-score gain under guidance (+11.7 points) is broadly distributed across Approach, Orchestration, Quality, and Feasibility, whereas GPT-5’s small net gain (+1.3 points) reflects offsetting movements: gains in Approach and Orchestration are partially cancelled by a Quality decrease, indicating that guidance helps GPT-5 discover the right tools but constrains its parameter choices. DeepSeek V3’s small net loss (−1.8 points) is concentrated in Quality and Feasibility, again consistent with guidance constraining an already-capable agent. These component-level patterns complement the stage-level heatmap by showing that guidance affects *which* aspects of the rubric improve, not just *whether* agents use more tools.

### De novo vs Redesign Performance Breakdown

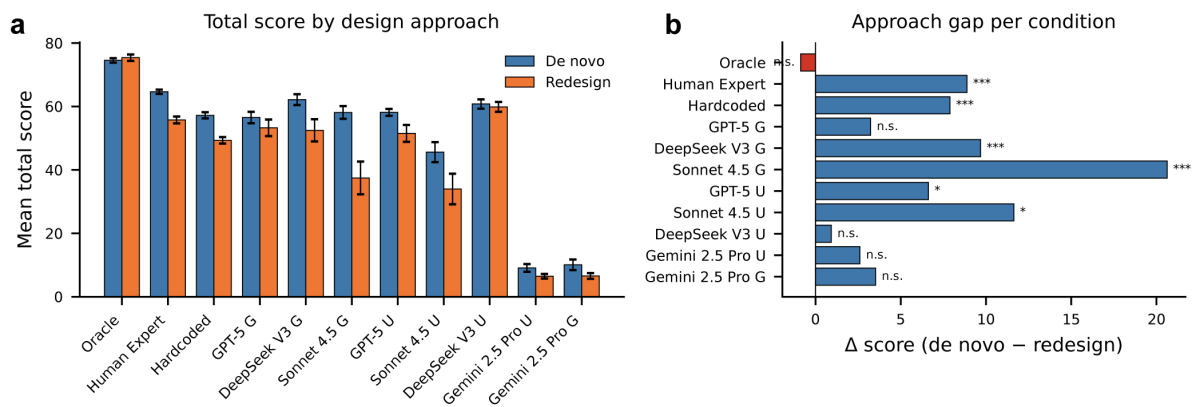

**Fig. S6 De novo versus redesign performance breakdown.** (a) Mean total score by design approach (*de novo* vs. redesign) for all 11 conditions, ordered by total score. Error bars show standard error of the mean. (b) Approach gap per condition, computed as $\Delta = \text{mean}(\text{de novo}) - \text{mean}(\text{redesign})$, with Mann–Whitney *U* significance annotations (\*$p < 0.05$, \*\*\*$p < 0.001$; n.s. = not significant). Oracle and Human Expert show the largest approach gaps; most LLM conditions show modest differences.

### Appendix K Variance Partitioning and Regression Analysis

This section provides full statistical details for the hierarchical regression analysis reported in the main text (Section 2.4).

Supplementary Figure S13 presents four complementary views of the variance-partitioning analysis. Panel (a) shows the variance partitioning as a stacked bar: binary coverage alone explains $R^2 = 0.514$; depth features add $+0.036$ ($p < 10^{-11}$); candidate features add $+0.012$ ($p = 0.001$); total $R^2 = 0.562$. When Gemini is excluded, binary coverage drops to $R^2 = 0.143$ and depth’s contribution increases dramatically to $+0.191$, confirming that depth is the primary differentiator among functional agents. The full nested-model comparison (with all conditions and excluding Gemini 2.5 Pro) achieves $R^2 = 0.545$ (all conditions) with the lowest AIC (7156.3) and BIC (7215.6).

Panel (b) shows LASSO feature importance: binary execution coverage is the strongest single predictor (coefficient = 11.39), and among depth features the number of distinct evaluation metric categories per candidate (coefficient = 3.57) persists longest, confirming that the *variety* of evaluation is a more robust predictor than raw evaluation frequency.

Panel (c) provides the component-level $R^2$ breakdown. Approach is almost entirely explained by binary features ($R^2 = 0.81$; depth lift negligible). Diversity shows the most dramatic depth dependence (binary $R^2 = 0.14$, depth lift $+0.186$). Feasibility shows a moderate depth lift ($+0.031$). Quality resists prediction from all feature sets ($R^2 < 0.16$), reflecting its dependence on biophysical properties not directly encoded in the trace features.

Panel (d) compares mean evaluations per candidate split by design approach for the expert and guided LLM conditions; the expert evaluates *de novo* candidates substantially deeper than redesign candidates while LLM agents show flat profiles — a depth-adaptation pattern explored in detail in Section M.

### Appendix L Generation versus Evaluation Decomposition

This section provides definitions of the depth metrics used throughout the manuscript and their per-condition values, decomposes the evaluation depth gap into candidate generation volume and per-candidate evaluation depth, and reports the candidate screening (filtering) behaviour referenced in the main text. All metrics are derived algorithmically from logged MCP tool invocations and require no subjective judgement.

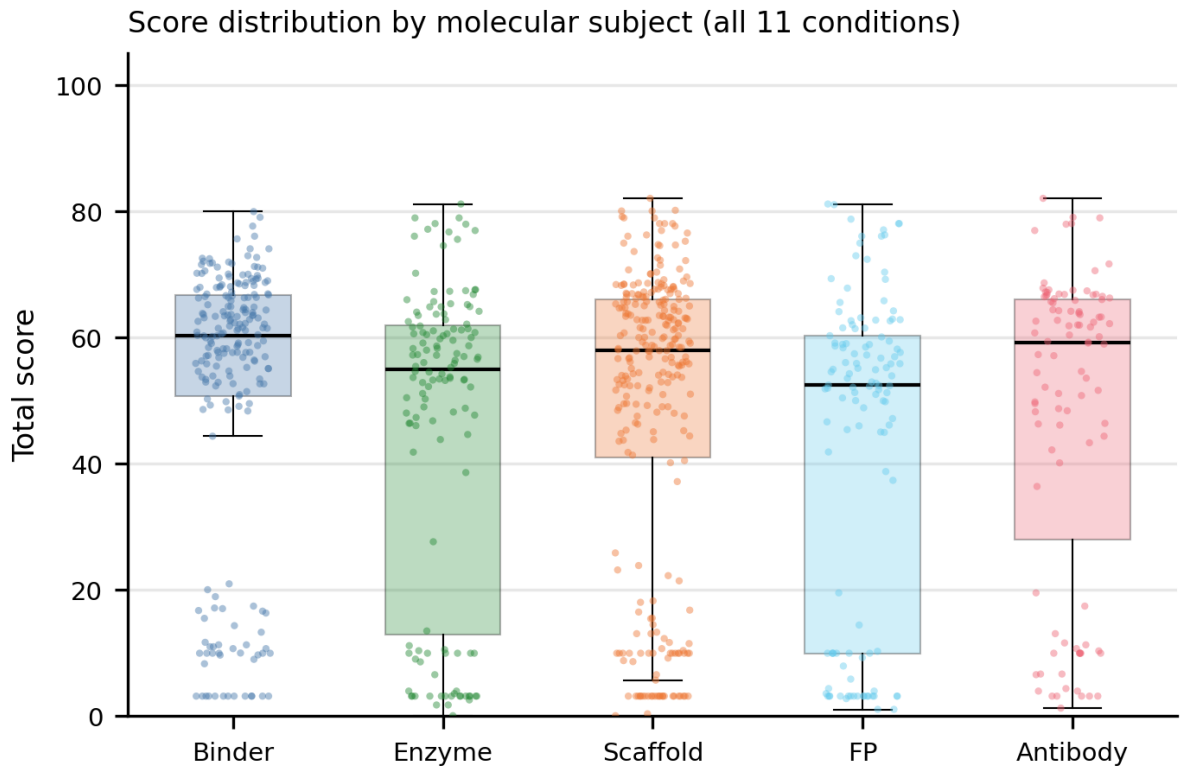

**Fig. S7 Score distribution by molecular subject across all 11 conditions.** Boxplots with overlaid strip plots of per-task total score, stratified by molecular subject (binder, enzyme, scaffold, fluorescent protein, antibody). Each panel shows the distribution for one of the 11 evaluation conditions (four LLMs × two modes, plus Hardcoded Pipeline, Human Expert, and Human Oracle). Binder tasks consistently yield the highest scores; antibody and fluorescent-protein tasks are the most challenging across conditions. The agent-expert gap is largest on antibody and fluorescent-protein tasks, foreshadowing the domain-specific depth analysis in Sections H–K.

### L.1 Depth metric definitions

We define three depth metrics. (i) The *evaluation-to-generation ratio* is the number of evaluative tool calls divided by the number of generative tool calls, where evaluative tools comprise structure prediction, stability scoring, interface analysis, energy minimization, and physics validation, and generative tools comprise backbone generation and sequence design. (ii) The *number of distinct evaluation metric categories per candidate* counts how many of six evaluation categories (structure prediction, interface analysis, physics-based scoring, binding assessment, stability analysis, sequence evaluation) are applied to each design candidate (range 0–6); this is the variety axis used in main-text Figs. 3d and 4d. (iii) The *tool-call rate per pipeline stage relative to the human expert* normalises per-stage tool counts against the human-expert reference, providing a stage-specific measure of utilisation intensity.

Across the 836 task-condition observations, the evaluation-to-generation ratio correlates strongly with total score ($\rho = 0.685$, $p < 10^{-117}$) and the number of distinct evaluation metric categories per candidate shows the largest expert-versus-LLM gap of any depth metric. The expert achieves a mean evaluation-to-generation ratio of 5.4 and a median of 4 distinct metric categories per candidate, while all LLM conditions remain in the ranges 1.0–2.0 and 1–2 respectively. Mann-Whitney $U$ tests confirm significant expert-vs-LLM differences for the scoring/validation stage in particular (all $p < 10^{-15}$), while backbone generation depth is statistically comparable between expert and the strongest LLM agents.

### L.2 Candidate generation versus per-candidate evaluation

Supplementary Figure S14 presents the generation-evaluation decomposition as a scatter plot of mean candidates per task versus mean evaluations per candidate (log scale) for all 11 conditions. The human expert occupies the upper-right quadrant with moderate candidate volume ($\sim 1.6$ candidates/task) but dramatically higher per-candidate evaluation depth ($5.9\times$ above the best LLM, all $p < 10^{-23}$, Cohen’s $d > 1.88$). LLM agents cluster in the lower portion of the plot with comparable candidate counts but substantially shallower evaluation. This decomposition confirms that the evaluation depth gap is driven entirely by per-candidate evaluation depth, not by candidate generation volume.

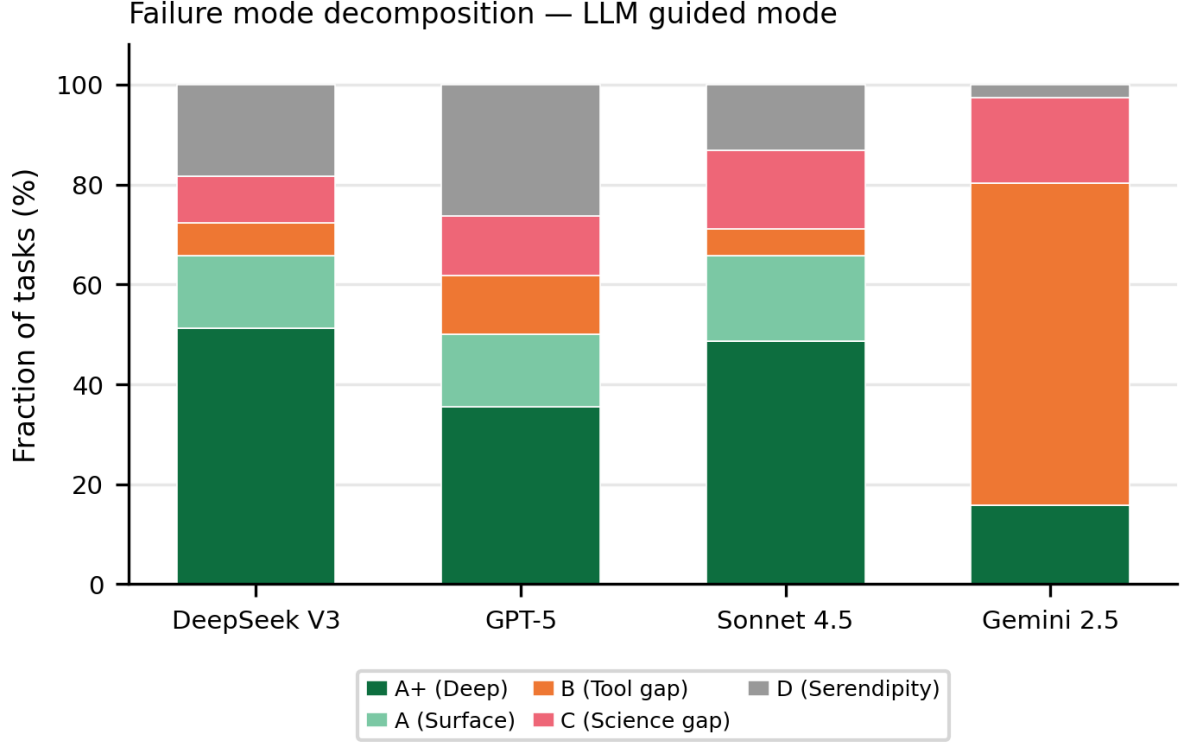

**Fig. S8 Failure mode decomposition for four LLM agents under guided mode.** Stacked bar chart showing the fraction of tasks in each of five failure-mode categories — deep success, shallow success, tool gap, science gap, serendipity — for DeepSeek V3, GPT-5, Claude Sonnet 4.5, and Gemini 2.5 Pro, all evaluated in guided mode across $n = 76$ tasks. DeepSeek V3 and GPT-5 are dominated by shallow success (correct plan, successful execution but shallow evaluation), with the largest deep-success share observed for DeepSeek V3. Sonnet 4.5 shows a larger science-gap share than the other top-tier models. Gemini 2.5 Pro is dominated almost entirely by science gap, reflecting its systematic MCP tool-calling failure. The serendipity category contributes negligibly across all models. Original $A^+/A/B/C/D$ legend retained in the figure for cross-reference; plain-English equivalents are used throughout the text.

#### L.3 Quality-gating and filtering

Complementary analyses of candidate screening behaviour confirm that no LLM agent condition exhibits a filter ratio significantly above $1.0\times$: all eight LLM conditions output the entirety of their generated candidate pool. The Hardcoded Pipeline achieves $1.7\times$, reflecting its built-in top-design selection. The Human Expert ($n = 1$) achieves $5.1\times$, generating a mean of 10.2 effective candidates per task and retaining 2.5; this value is reported as a single-expert reference. Separately, the number of effective candidates shows no significant correlation with the Diversity rubric component ($\rho = 0.055$, $p = 0.11$), while the number of independent backbone generation calls strongly predicts Diversity ($\rho = 0.576$, $p < 10^{-50}$).

#### L.4 Stochastic repetition index

To assess whether agents differentially repeat stochastic versus deterministic tools, we computed the Stochastic Repetition Index (SRI), defined as the mean number of calls per unique stochastic tool type divided by the mean number of calls per unique deterministic tool type. $SRI > 1$ indicates generation-dominant usage; $SRI < 1$ indicates evaluation-dominant usage. Counter to an initial hypothesis that expert practitioners would preferentially repeat stochastic tools, the Expert achieves a mean SRI of 0.62, lower than all LLM agents (pooled LLM mean = 1.52; Mann–Whitney $p = 7.67 \times 10^{-8}$, Cohen’s $d = -0.65$; Supplementary Table S2). This reversal reflects the Expert’s evaluation-dominant strategy: 26.4 deterministic evaluation calls per task versus 4.9 stochastic generation calls. LLM agents’ $SRI > 1$ arises from shallow evaluation (3.8–5.2 deterministic calls) combined with moderate generation (3.0–5.5 stochastic calls).

### Appendix M Depth Adaptation by Task Type

Supplementary Figure S15 examines how depth adaptation differs between *de novo* and redesign tasks. The expert systematically adjusts evaluation depth between approaches — deeper evaluation on *de novo* tasks (mean evaluation-to-generation ratio $\approx 25$) than on redesign tasks ($\approx 5$) — whereas all LLM conditions show flat depth profiles across

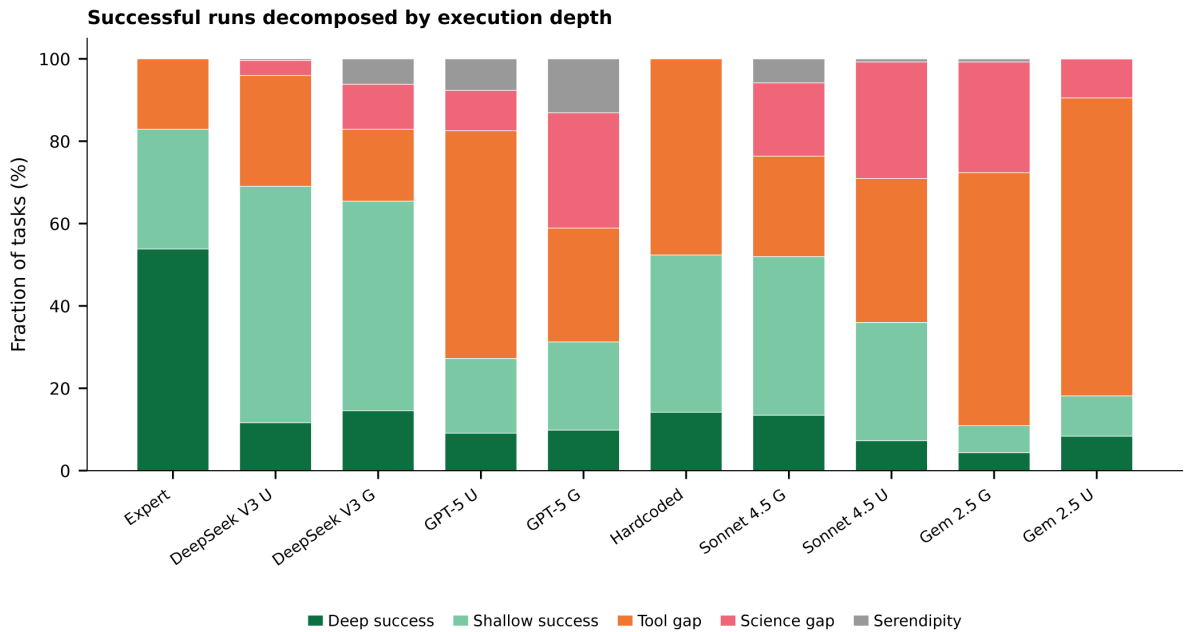

**Fig. S9 Successful runs decomposed by execution depth across all 11 conditions.** Stacked bar chart showing, for every evaluation condition, the fraction of tasks falling into deep success (above expert-median execution depth), shallow success (below expert-median execution depth), tool gap, science gap, and serendipity. The human expert achieves deep success on 53.8% of tasks; the strongest LLM (DeepSeek V3 unguided) reaches deep success on only 14.5% — a $3.7\times$ gap that is invisible in binary success/failure analysis. Hardcoded pipeline shows a moderate deep-success rate from its built-in evaluation loop. Gemini conditions are dominated by tool gap, consistent with their MCP tool-calling failure.

**Table S2** Stochastic Repetition Index (SRI) across evaluation conditions. SRI $< 1$ indicates evaluation-dominant usage. $p$: two-sided Mann–Whitney $U$ vs. Human Expert; $d$: Cohen’s $d$.

| Condition | SRI | Stoch. calls | Det. calls | $p$ | $d$ |
| --- | --- | --- | --- | --- | --- |
| Human Expert | 0.62 | 4.9 | 26.4 | — | — |
| DeepSeek V3 guided | 1.12 | 2.9 | 5.7 | $3.87 \times 10^{-4}$ | -0.37 |
| DeepSeek V3 unguided | 1.51 | 4.5 | 5.2 | $3.20 \times 10^{-8}$ | -1.02 |
| GPT-5 guided | 1.66 | 3.3 | 3.8 | $1.67 \times 10^{-7}$ | -0.77 |
| GPT-5 unguided | 2.82 | 5.5 | 5.1 | $2.18 \times 10^{-11}$ | -1.02 |
| Sonnet 4.5 guided | 1.19 | 3.5 | 4.2 | $4.30 \times 10^{-4}$ | -0.68 |
| Sonnet 4.5 unguided | 1.25 | 3.5 | 3.8 | $9.25 \times 10^{-1}$ | -0.45 |
| Hardcoded Pipeline | 1.26 | 2.2 | 3.7 | $8.61 \times 10^{-5}$ | -0.69 |

approaches (mean differences $< 0.5$). This depth-adaptation gap is most pronounced in the scoring and refinement stages and is consistent with the broader finding that LLM agents apply a single shallow evaluation template regardless of task type.

### Appendix N Forced-Depth Intervention Design and Subset Validation

This section describes the design and execution of the forced-depth intervention experiments reported in the main text (Section 2.5).

#### N.1 Stratified subset selection

Because intervention runs are computationally expensive (each task–condition pair requires a fresh agent session with extended evaluation depth, increasing wall-clock time by $1.5\text{--}2.5\times$ relative to the baseline), we evaluated the intervention on a stratified subset of 18 tasks drawn from the full 76-task benchmark. The subset was constructed to preserve the joint distribution of design approach, molecular subject, and difficulty: 11 *de novo* and 7 redesign tasks; 4 antibody, 3 enzyme, 4 binder, 5 scaffold, and 2 fluorescent-protein tasks; 5 easy, 7 medium, and 6 hard tasks. This ensures that the intervention sample is representative of the full benchmark along all three stratification axes.

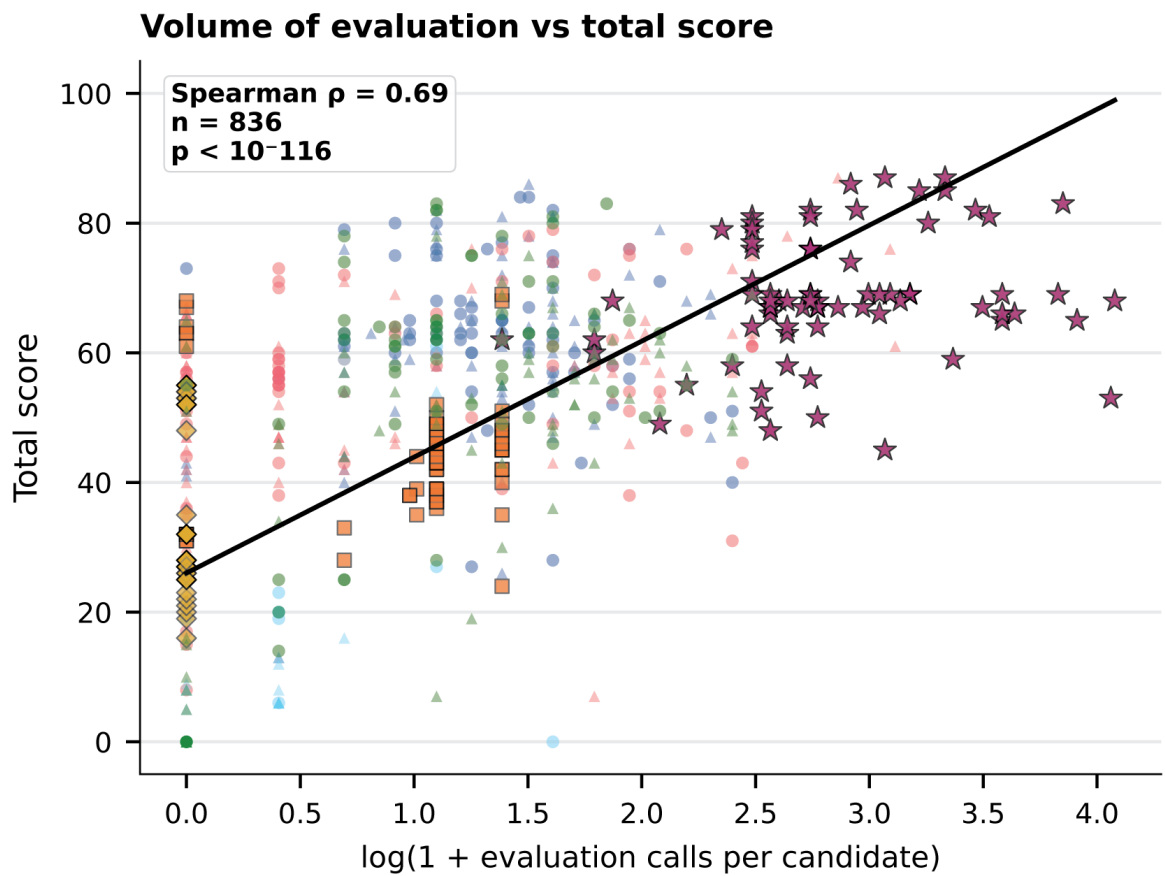

**Fig. S10 Volume of evaluation per candidate predicts total score.** Scatter of $\log(1 + \text{number of evaluation tool calls per candidate})$ against total score across all 836 task-condition observations (Spearman $\rho = 0.69$, $p < 10^{-116}$). Marker shape and colour identify the evaluation condition (DeepSeek V3, GPT-5, Sonnet 4.5, Gemini 2.5 Pro, Hardcoded, Human Expert; guided/unguided). Black line: linear fit. This is the volume-axis counterpart to main-text Fig. 3d, which uses the variety axis (number of distinct evaluation metric categories per candidate).

To verify representativeness, we compared per-task baseline scores between the 18-task subset and the full 76-task benchmark using two-sided Mann–Whitney $U$ tests for each condition, supplemented by category-level $\chi^2$ tests for taxonomy and difficulty distributions. Supplementary Figure S16 reports the results: all condition-level Mann–Whitney $p$-values exceed 0.30, the taxonomy $\chi^2$ test is non-significant ($p = 0.71$), and the difficulty $\chi^2$ test is non-significant ($p = 0.58$), confirming that the subset is statistically indistinguishable from the full benchmark on all measured axes. Median baseline scores on the subset are within $\pm 1.8$ points of the full-benchmark medians for all conditions.

### N.2 Forced-depth prompt template

The forced-depth intervention modifies only the agent’s system prompt; all MCP tool definitions, task prompts, and execution infrastructure are held constant relative to the baseline guided-mode condition. The intervention prompt appends a structured depth requirement to the baseline system prompt:

[Baseline system prompt ...]

REQUIRED EVALUATION PROTOCOL. For every candidate sequence you generate, you MUST execute at least the following evaluation steps before producing your final answer:

1. Predict the structure of the candidate using `predict_structure` (or `predict_complex` for binding tasks).
2. Score the predicted structure with at least *three* of the following tool categories: stability scoring, interface analysis, energy minimization, physics validation, sequence scoring.
3. Inspect the per-residue confidence (pLDDT) and any interface metrics (ipTM, $\Delta\Delta G$) returned by the evaluation tools.

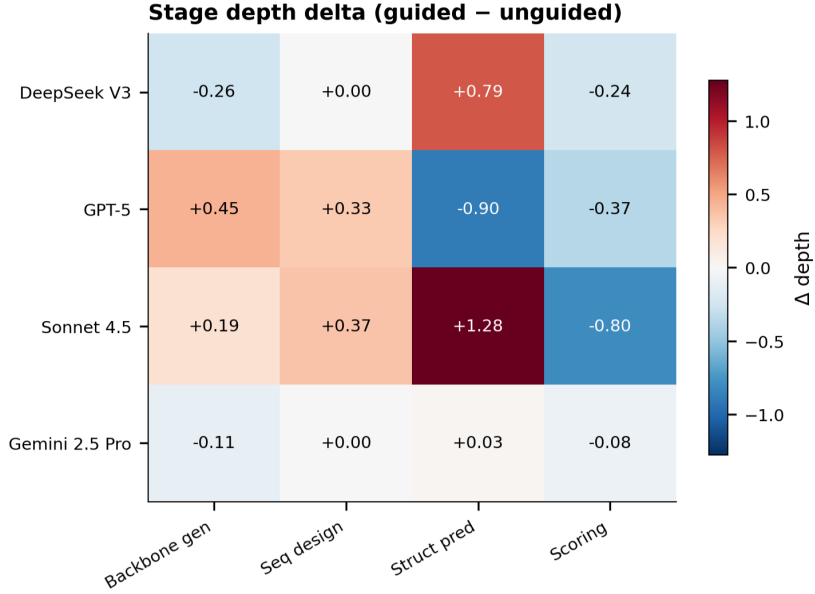

**Fig. S11 Stage depth delta between guided and unguided modes.** Heatmap of $\Delta\text{depth} = \text{guided} - \text{unguided}$ for each of four pipeline stages (backbone generation, sequence design, structure prediction, scoring) across the four LLM families (DeepSeek V3, GPT-5, Sonnet 4.5, Gemini 2.5 Pro). Positive values (red) indicate higher per-task stage depth in guided mode; negative values (blue) indicate higher depth in unguided mode. Sonnet 4.5 shows the largest guided-mode gain in structure prediction (+1.28), while GPT-5 shows a guided-mode decrease in structure prediction (−0.90), suggesting model-specific responses to enhanced tool descriptions.

4. If any evaluation metric falls below the documented thresholds, perform at least one round of refinement (re-design, re-optimize, or generate an additional candidate) before producing the final answer.

You may not skip these steps. Do not produce a final answer until the evaluation protocol has been completed for every candidate.

This protocol is designed to elevate the number of distinct evaluation metric categories applied per candidate from the LLM baseline range (1–2) towards the human-expert range (3–5), without prescribing any specific tool or any specific generative strategy. Critically, the prompt does not specify which tools to use, only that at least three distinct evaluation categories must be exercised — the agent retains discretion over tool selection, parameters, and ordering.

#### N.3 Loop budget and termination

To prevent runaway tool-call sequences while permitting the elevated evaluation depth, intervention runs use a per-task tool-call budget of 40 (versus 20 for baseline runs) and a wall-clock timeout of 300 s (versus 120 s for baseline). Agents that complete the protocol before exhausting their budget terminate normally; agents that exceed either limit have their session terminated and their final answer scored on whatever evaluation has been completed up to that point. We log the termination cause for each run and report distributions in Section T.

### Appendix O Within-Task Paired Comparison

Supplementary Figure S17 shows the within-task paired comparison for the intervention. DeepSeek V3 improves on 14/18 tasks (78% win rate); GPT-5 improves on 15/18 tasks (83%). The two failures-to-improve in DeepSeek and the three in GPT-5 cluster among tasks with already-high baseline scores, consistent with diminishing returns once baseline evaluation depth is sufficient.

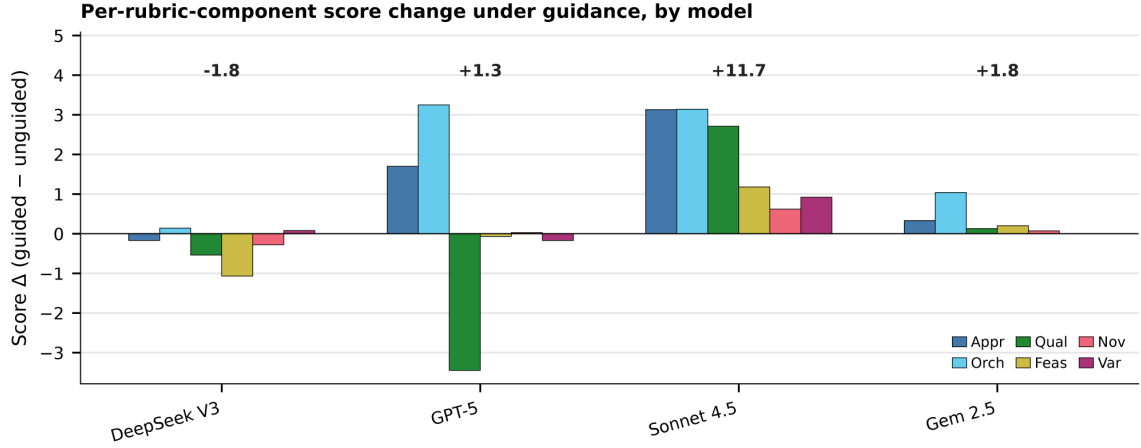

**Fig. S12 Per-rubric-component score change under guidance, by model.** Score delta (guided – unguided) decomposed into the six rubric components — Approach (Appr), Orchestration (Orch), Quality (Qual), Feasibility (Feas), Novelty (Nov), Diversity (Var) — for each LLM family. Numbers above the bars indicate the total delta summed across components. Sonnet 4.5 shows the largest broadly distributed gain (+11.7); GPT-5 (+1.3) and Gemini 2.5 Pro (+1.8) show small net effects with offsetting component changes; DeepSeek V3 (–1.8) is slightly harmed by guidance, consistent with prescriptive workflows constraining an already-capable agent.

### Appendix P Per-Rubric-Component Breakdown of Intervention

Supplementary Figure S18 provides the full per-rubric-component breakdown across all three intervention conditions (baseline, low-variety, forced-depth) for both models. The forced-depth gain is concentrated in Approach (and Orchestration for GPT-5) while Quality scores remain stable across conditions, confirming that the intervention changes how agents orchestrate tools rather than the biophysical output of the tools themselves. This pattern is consistent with the variance partitioning result (Section K) showing that Quality resists prediction from orchestration features.

### Appendix Q Manipulation Check

Supplementary Figure S19 confirms that the prompt manipulation produced its intended effect on the manipulation variable. The baseline mean number of distinct evaluation metric categories per candidate is 3.00 for DeepSeek V3 and 1.33 for GPT-5; the low-variety condition keeps these values close to baseline (3.33 and 1.61), while forced-depth raises them to 4.06 and 4.27. The two intervention conditions thus differ along the intended axis (variety) while sharing a similar tool-call budget, providing the experimental contrast that isolates evaluation variety as the operative factor in main-text Fig. 4.

### Appendix R Volume- versus Variety-Limited Models

Supplementary Figure S20 plots, for each model, the score gain achievable from added evaluation effort alone (low-variety – baseline) against the model’s baseline number of distinct metric categories per candidate. DeepSeek V3 occupies the variety-limited region of this plot: its baseline already applies three categories per candidate, and adding effort without adding variety yields essentially no score gain. GPT-5 occupies the volume-limited region: its baseline applies fewer than two categories per candidate, and any added effort yields large gains regardless of whether new categories are introduced. This model-dependent split predicts that as future LLMs raise their default evaluation effort, structured variety will become the dominant lever.

### Appendix S Baseline Depth Profile and Intervention Response

This section reports the per-task and per-condition outcomes of the forced-depth intervention summarised in the main text (Section 2.5).

The two LLMs evaluated under the intervention — DeepSeek V3 and GPT-5, both in guided mode — show large and statistically significant depth elevation and corresponding score gains. DeepSeek V3 mean variety (number of distinct evaluation metric categories per candidate) rises from 3.00 (baseline) to 4.06 (intervention), and mean total score rises from 58.7 to 68.1 (Wilcoxon signed-rank $p = 0.002$, Cohen’s $d = 0.58$, $n = 18$). GPT-5 mean variety rises from 1.33 to 4.27, and mean total score rises from 46.8 to 62.7 (Wilcoxon signed-rank $p < 0.001$, Cohen’s $d = 1.27$, $n = 18$). Both improvements survive Bonferroni correction at $\alpha = 0.0083$.

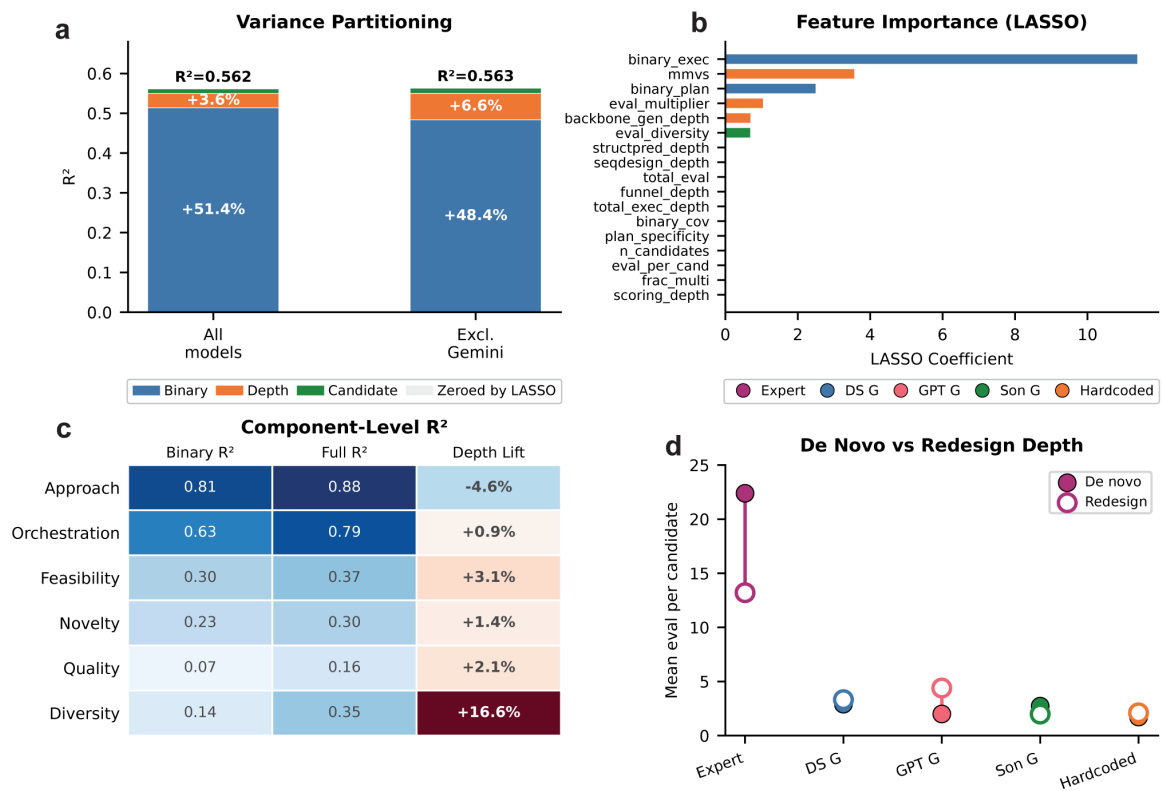

**Fig. S13 Component-level $R^2$ and variety-of-evaluation lift decomposition.** (a) Variance partitioning stacked bar. Left: all conditions ($R^2 = 0.562$; binary 51.4%, depth +3.6%, candidate +1.2%). Right: excluding Gemini ($R^2 = 0.563$; binary 48.4%, depth +6.6%). Depth’s relative contribution increases when the Gemini floor effect is removed. (b) LASSO feature importance. Binary execution coverage is the strongest predictor; among depth features the number of distinct evaluation metric categories per candidate persists longest, confirming variety over raw frequency. (c) Component-level $R^2$ table for the six rubric components. Columns show binary-only $R^2$, full-model $R^2$, and depth lift ($\Delta R^2$). Approach is almost entirely explained by binary features ($R^2 = 0.81$); Diversity shows the most dramatic depth dependence (binary $R^2 = 0.14$, depth lift +16.6%); Quality resists prediction ($R^2 < 0.16$). (d) *De novo* versus redesign evaluation depth. Mean evaluations per candidate for expert and guided LLM conditions, split by approach. The expert evaluates *de novo* candidates at substantially higher depth than redesign candidates; LLM agents show flat profiles across approaches.

Supplementary Figure S21 characterises the baseline depth profiles and intervention response for both models. GPT-5 starts from a substantially lower baseline (mean of 1.33 distinct metric categories per candidate) and fewer evaluation calls per task (3.0) compared to DeepSeek V3 (3.00 categories, 5.6 calls/task), explaining its larger intervention effect. The forced-depth intervention raises the variety axis for both models while the low-variety control does not, and component-level score deltas reveal that DeepSeek V3 gains broadly across all rubric components whereas GPT-5’s gains are concentrated in Approach. Per-task data tables reporting baseline score, intervention score, $\Delta$, and termination cause for all 18 tasks × 2 conditions are provided in the auxiliary supplementary spreadsheet.

### Appendix T Termination Behaviour Under Intervention

Supplementary Figure S22 reports the distribution of termination causes (normal completion, tool-call budget exhaustion, wall-clock timeout) for baseline and intervention runs. Under the intervention, 14/18 DeepSeek V3 and 15/18 GPT-5 runs terminate normally; the remainder hit the tool-call budget but still complete the required evaluation protocol before producing a final answer. The cumulative distribution of evaluation calls per task shifts rightward under forced-depth for both models, while the rate of immediate submissions (zero evaluation calls before submission) drops from $\sim 33\%$ for GPT-5 baseline to 0% under forced-depth, indicating that agents respect the protocol minimum.

### Appendix U Candidate Quality Distributions Under Intervention

Supplementary Figure S23 compares per-candidate quality distributions between baseline and intervention runs. Intervention runs do not generate qualitatively different candidates — the distribution of pTM and ipTM across the candidate pool is statistically indistinguishable from baseline (Kolmogorov–Smirnov $p > 0.30$ for both metrics in both models). What changes is selection: under the intervention, agents systematically retain higher-quality candidates from a similar candidate pool, consistent with a screening-driven mechanism rather than a generation-driven one. Together with the manipulation check (Section Q), termination behaviour (Section T), and the volume–variety regime analysis (Section R), this supports the interpretation that forced-depth gains arise from the intended mechanism — elevated per-candidate evaluation variety and selection pressure — and not from artifacts such as runtime inflation or candidate-count escalation.

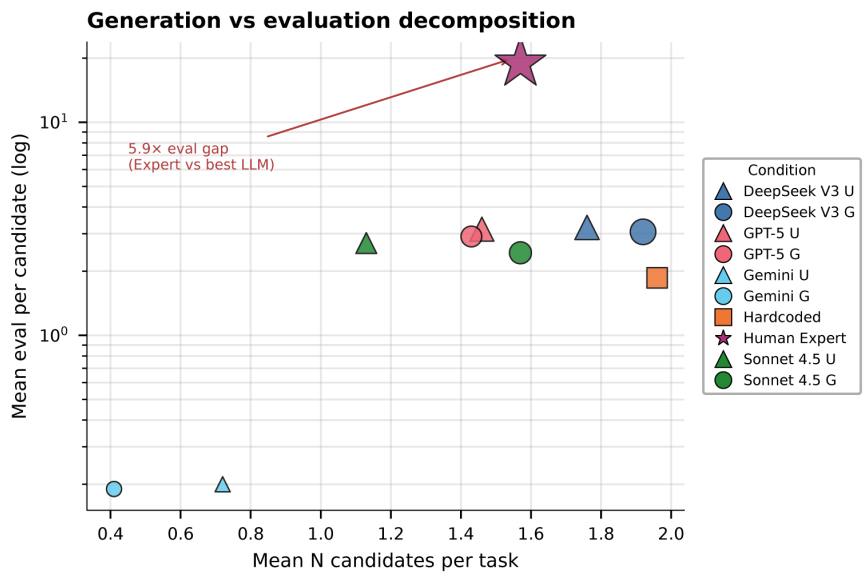

**Fig. S14 Generation versus evaluation decomposition.** Scatter plot of mean candidates generated per task ($x$-axis) versus mean evaluations per candidate ($y$-axis, log scale) for all 11 conditions. Marker shapes distinguish guided (triangle) and unguided (circle) modes; colours indicate LLM family. The human expert (star) occupies the upper-right quadrant with moderate candidate volume ($\sim 1.6$ candidates/task) but dramatically higher per-candidate evaluation depth ($5.9\times$ above the best LLM agent), confirming that the evaluation depth gap is driven by per-candidate depth rather than candidate generation volume. LLM agents cluster in the lower portion of the plot, with DeepSeek V3 achieving the highest candidate count among LLMs.

### Appendix V De novo versus Redesign Intervention Effect

Supplementary Figure S24 reports mean Δscore (forced-depth – baseline, ± SEM) split by design approach. The intervention effect is positive on both *de novo* and redesign tasks for both models, with modestly larger effects on *de novo* tasks, mirroring the larger baseline depth-adaptation gap on generative tasks reported in Section M. This subgroup pattern provides a second, independent line of evidence that the intervention closes the specific behavioural gap identified in the diagnostic analyses.

### Appendix W Domain-Specific Intervention Heterogeneity

Supplementary Figure S25 reports per-task Δscore (forced-depth – baseline) by molecular-subject domain for both models. Due to the stratified subset design ($n = 2\text{--}4$ tasks per domain–model cell), these results are descriptive rather than inferential. Both models show positive effects across most domains. GPT-5 shows the largest gains in scaffold and fluorescent-protein domains; DeepSeek V3 shows the largest gains in scaffold. This pattern is broadly consistent with the interpretation that the intervention closes a depth deficit, with domain-level variation reflecting differences in baseline depth gap magnitude.

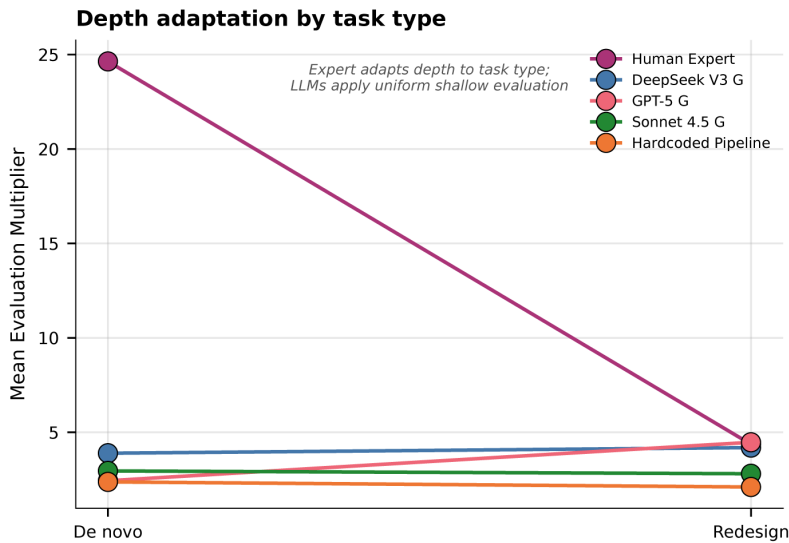

**Fig. S15 Depth adaptation by task type.** Mean evaluation-to-generation ratio for the human expert (red), DeepSeek V3 guided (blue), GPT-5 guided (green), Sonnet 4.5 guided (orange), and the hardcoded pipeline (pink) on *de novo* versus redesign tasks. The expert dramatically reduces evaluation effort on redesign tasks (from $\sim 25\times$ to $\sim 5\times$), demonstrating adaptive depth allocation. All LLM agents and the hardcoded pipeline show flat profiles ($\approx 3\text{--}5$) regardless of approach, consistent with a single shallow evaluation template applied uniformly across task types.

### Appendix X Robustness of Central Findings

Three potential concerns about the generality of the central findings are: (1) Gemini 2.5 Pro’s floor-level performance may inflate statistical relationships; (2) the single human expert ($n = 1$) may disproportionately anchor depth–score correlations; and (3) the intervention effect may be driven by a single molecular-subject domain. We address all three through systematic exclusion analyses (Supplementary Figure S26).

#### X.1 Gemini 2.5 Pro exclusion

The coverage–depth dissociation reported in the main text is robust to Gemini exclusion. The unguided-to-guided change in coverage and in depth, normalised to the human-expert ceiling and recomputed for each LLM family with and without Gemini, places Sonnet, GPT-5, and DeepSeek along the coverage axis with near-zero movement on the depth axis in both versions, confirming that the dissociation is not an artifact of Gemini’s tool-calling failure. Variance partitioning likewise tells a stronger story when Gemini is excluded: the variety axis’s $R^2$ contribution nearly doubles from 4% ($p = 9 \times 10^{-11}$) to 7% ($p < 10^{-16}$), reflecting that Gemini’s universal floor effect compresses the contribution of any continuous predictor.

#### X.2 Expert exclusion

The variety-to-score correlation persists when expert observations are removed. Across the 836 task–condition observations, the full-sample Spearman $\rho$ is 0.685 ($p < 10^{-117}$); restricting to LLM-only conditions yields $\rho = 0.68$, comparable to the full sample. This confirms that the depth–score relationship is established within the LLM population and is not an artifact of a single-expert anchor.

#### X.3 Domain leave-one-out

The forced-depth intervention effect is robust to single-domain exclusion. Recomputing Cohen’s $d$ on the 18-task subset with each molecular-subject domain removed in turn yields effect sizes within $\pm 0.15$ of the full-subset values for both DeepSeek V3 and GPT-5, with all leave-one-out effects remaining large ($d > 0.4$ for DeepSeek; $d > 1.0$ for GPT-5). No single domain drives the intervention result.

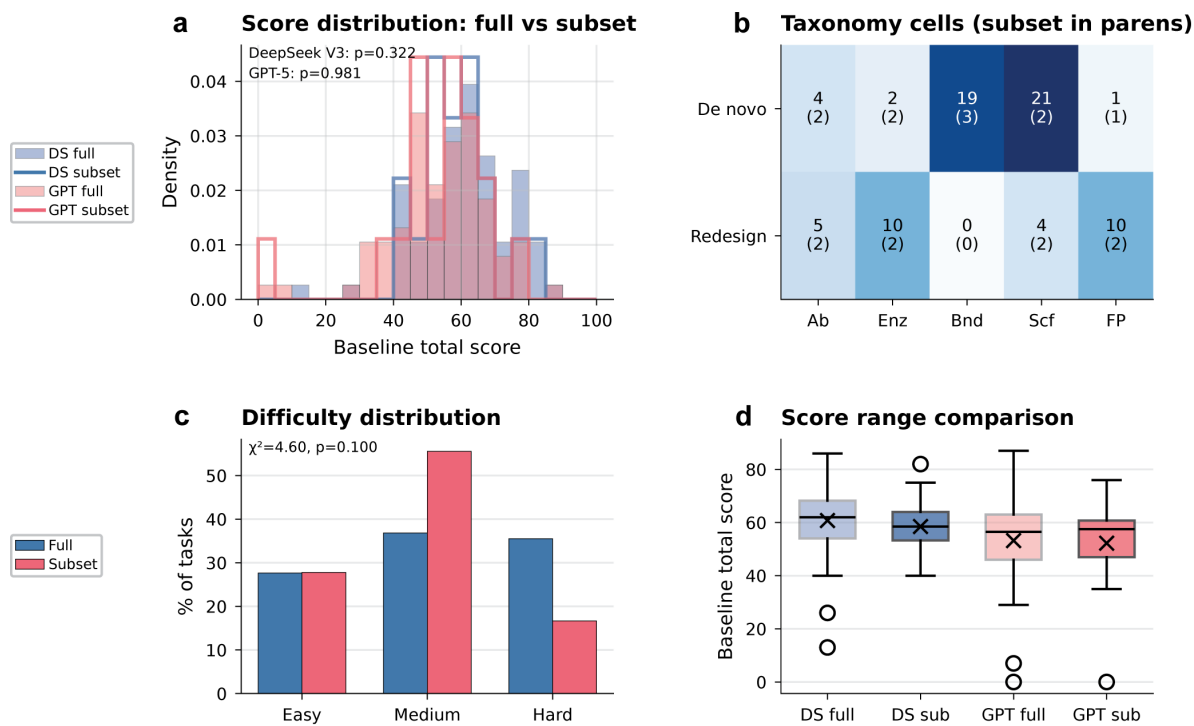

**Fig. S16 Subset representativeness validation.** (a) Baseline total score density distributions for the full benchmark (solid) and the 18-task intervention subset (dashed) for DeepSeek V3 and GPT-5. Mann–Whitney $U$ tests confirm no detectable distributional shift (DeepSeek $p = 0.322$; GPT-5 $p = 0.981$). (b) Taxonomy cell counts for the full $2 \times 5$ approach × molecular-subject matrix, with subset counts in parentheses. The subset preserves the joint distribution across all populated cells. (c) Difficulty distribution for the full benchmark (blue) versus the subset (red); $\chi^2$ test $p = 0.100$, confirming no significant difficulty imbalance. (d) Score range comparison (boxplot) for DeepSeek V3 and GPT-5 on the full benchmark versus the subset, showing overlapping distributions and comparable medians.

Supplementary Figure S26 summarises these robustness checks in a single $2 \times 2$ panel layout combining the coverage–depth dissociation, depth–score correlation, variance partitioning, and intervention effect size, each shown with and without the relevant exclusion. All four central findings remain qualitatively unchanged.

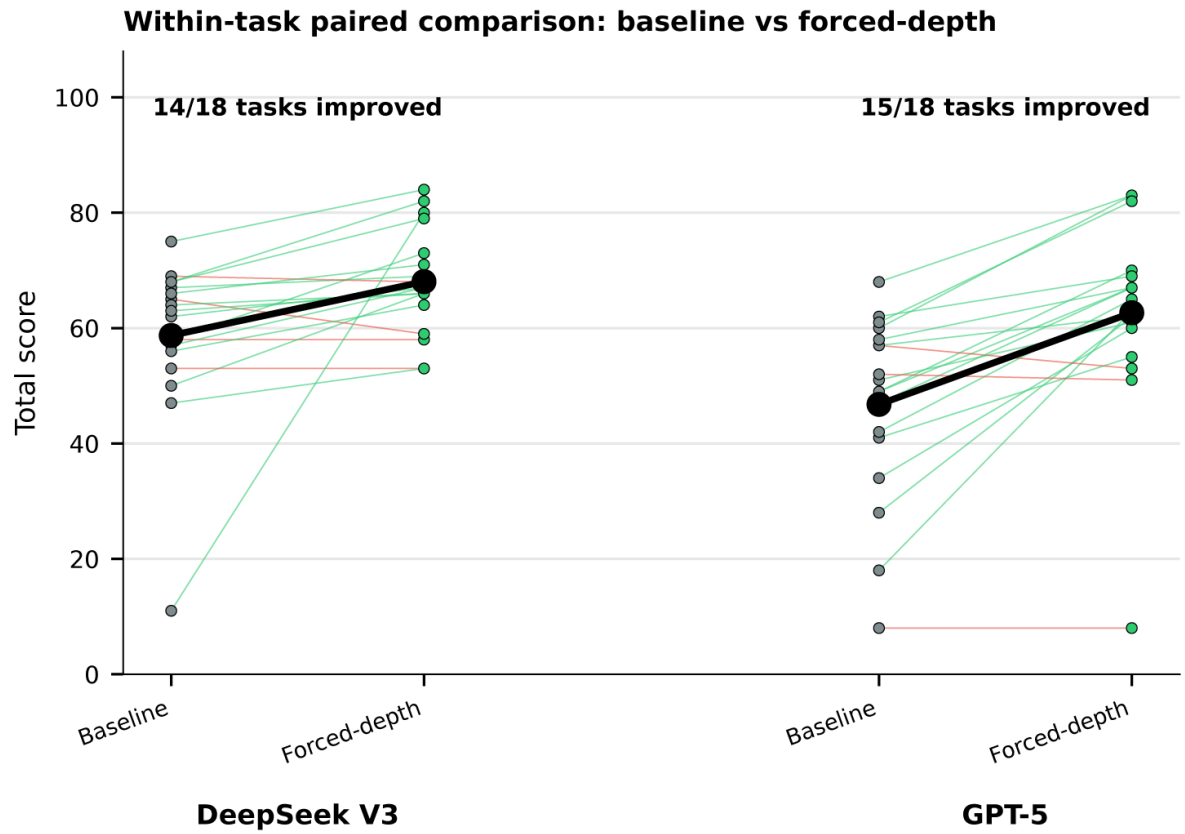

**Fig. S17 Within-task paired comparison: baseline versus forced-depth.** Total score for each of the 18 tasks under baseline (grey) and forced-depth (green), connected by lines (green if improved, red if worsened) for DeepSeek V3 (left) and GPT-5 (right). Bold black points and line indicate condition means. DeepSeek V3 improves on 14/18 tasks (78% win rate); GPT-5 improves on 15/18 tasks (83%). The two failures-to-improve in DeepSeek and the three in GPT-5 cluster among tasks with already-high baseline scores, consistent with diminishing returns once baseline evaluation depth is sufficient.

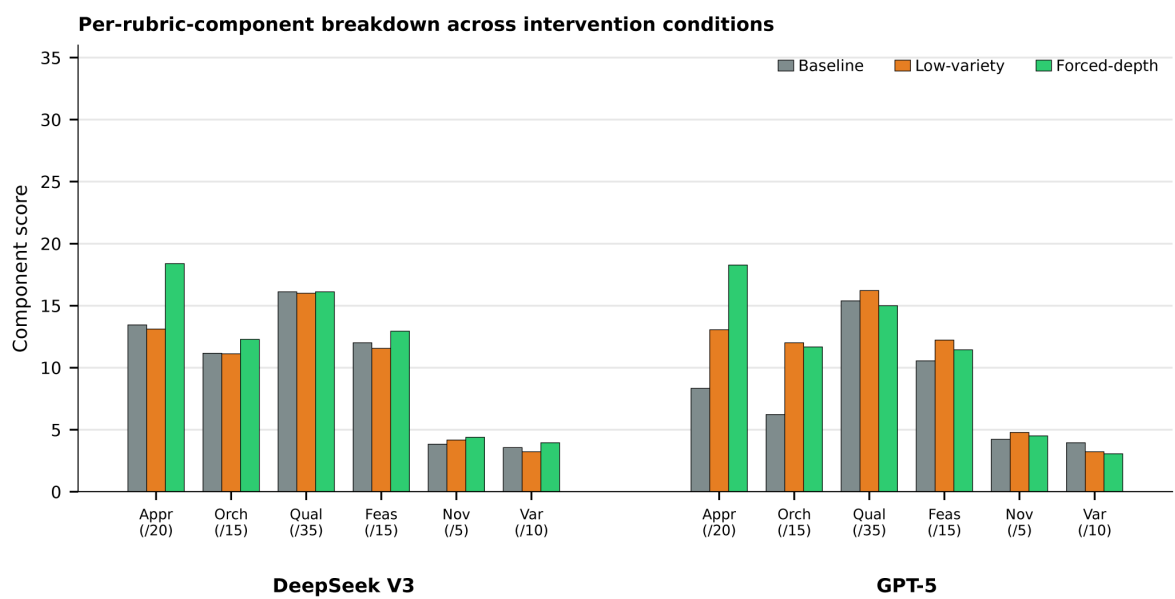

**Fig. S18 Per-rubric-component breakdown across intervention conditions.** Component scores (out of the rubric maximum for each component: Approach 20, Orchestration 15, Quality 35, Feasibility 15, Novelty 5, Diversity 10) under three conditions (baseline grey; low-variety orange; forced-depth green) for DeepSeek V3 (left) and GPT-5 (right) on the 18-task subset. The forced-depth gain is concentrated in Approach (and Orchestration for GPT-5), while Quality scores remain stable across conditions, confirming that the intervention changes how agents orchestrate tools rather than the biophysical output of the tools themselves.

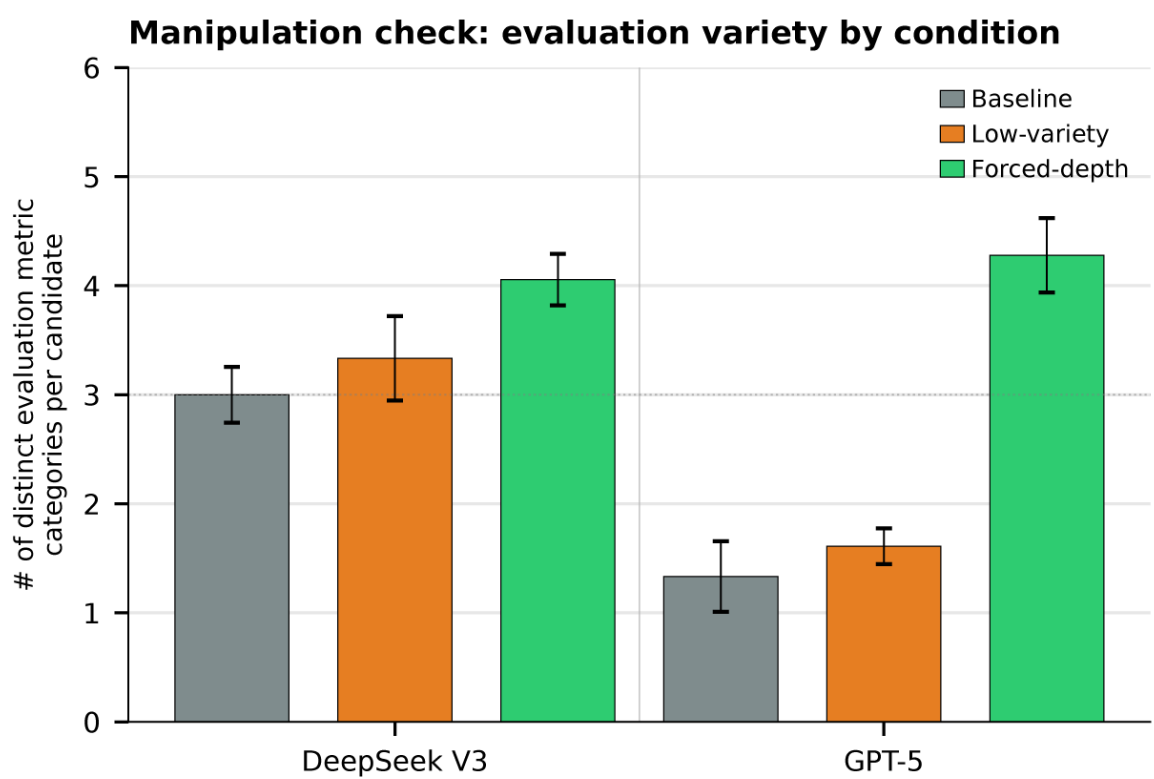

**Fig. S19 Manipulation check: number of distinct evaluation metric categories per candidate by condition.** Bar chart of the mean number of distinct evaluation metric categories per candidate (out of six) for DeepSeek V3 and GPT-5 under three conditions (baseline grey, low-variety orange, forced-depth green). Forced-depth raises this number substantially in both models, while low-variety remains close to baseline. The two intervention conditions therefore differ along the intended axis (variety of evaluation), enabling the matched-compute contrast in main-text Fig. 4 to isolate variety as the operative factor.

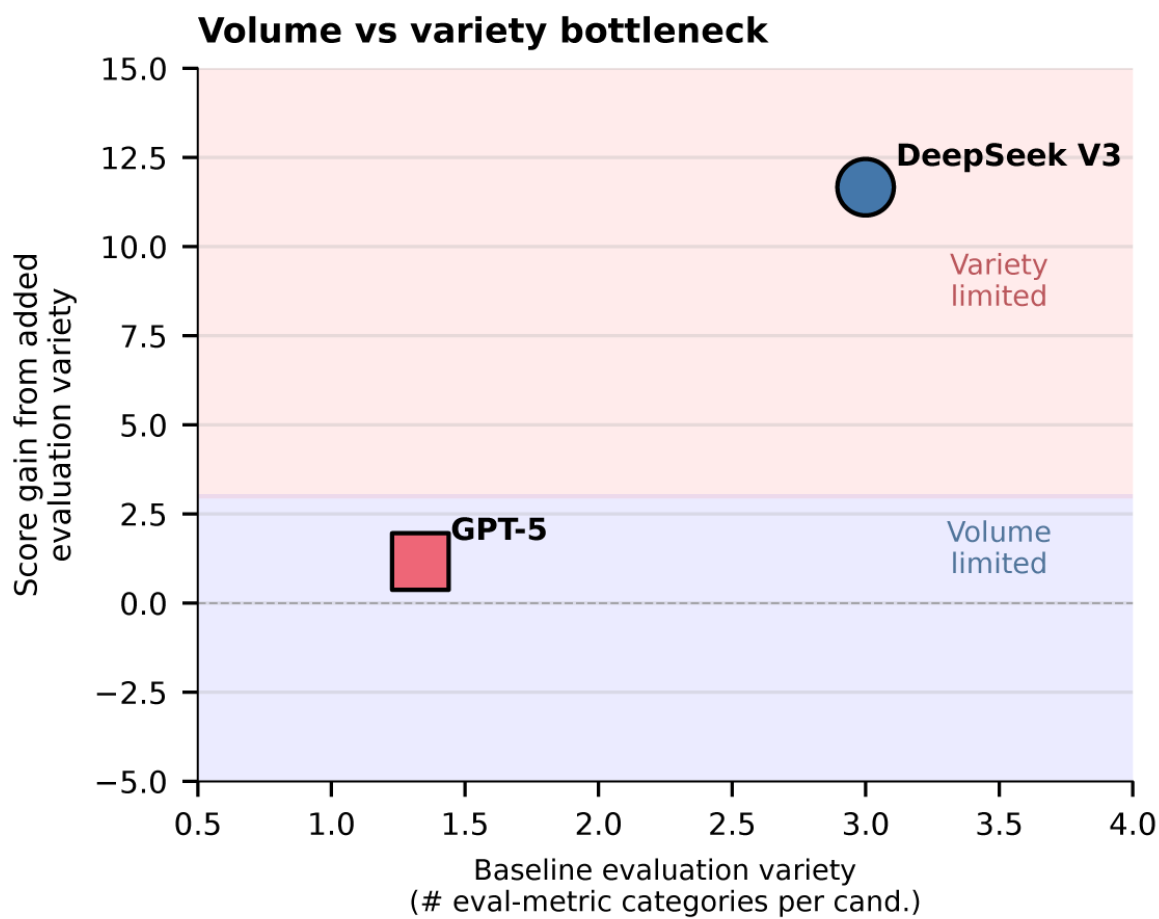

**Fig. S20 Volume- versus variety-limited models.** Score gain from added evaluation effort alone (low-variety – baseline) plotted against the baseline number of distinct evaluation metric categories per candidate, for DeepSeek V3 and GPT-5. DeepSeek V3 (top right) is variety-limited: its baseline already applies $\sim 3$ categories per candidate, so adding effort without adding variety yields negligible gain. GPT-5 (bottom centre) is volume-limited: its baseline applies fewer than two categories per candidate, so any added effort yields large gains regardless of whether new categories are introduced. The shaded regions indicate the qualitative regimes; the model-dependent placement predicts that as future LLMs raise default evaluation effort, the marginal value of structured variety will increase.

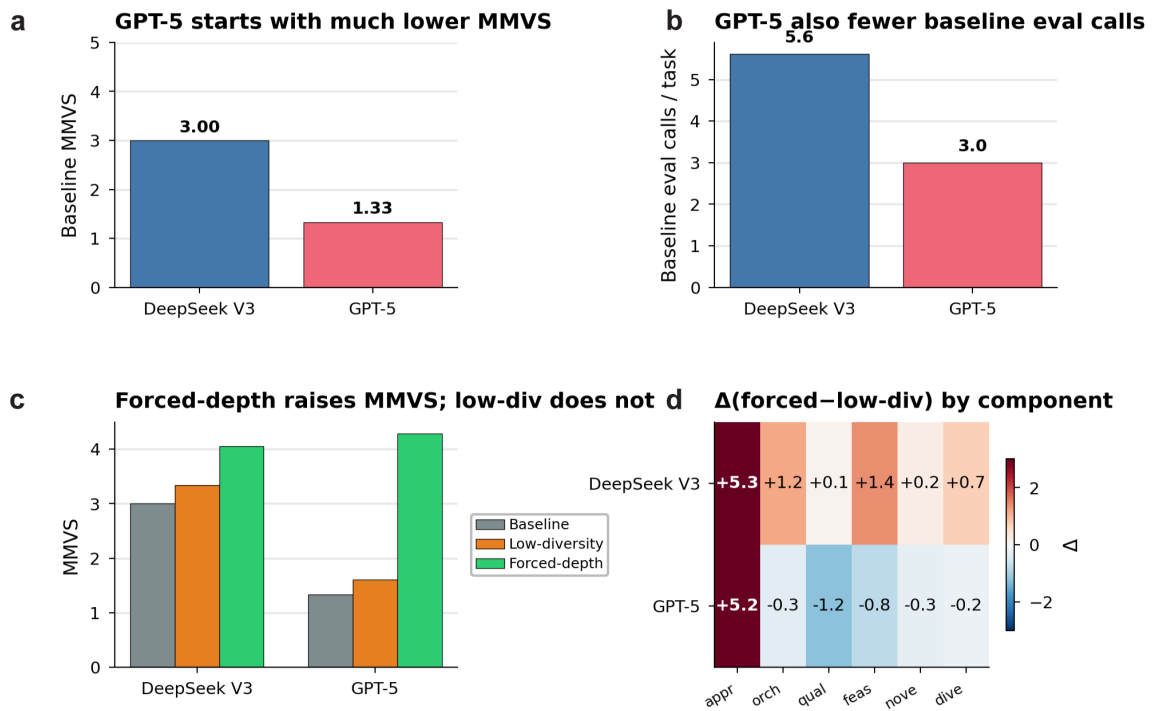

**Fig. S21 GPT-5 versus DeepSeek V3 baseline depth profile and intervention response.** (a) Baseline variety comparison — GPT-5 starts with substantially fewer distinct evaluation metric categories per candidate (1.33) than DeepSeek V3 (3.00), indicating shallower multi-metric evaluation at baseline. (b) Baseline evaluation call volume — GPT-5 also makes fewer evaluation calls per task (3.0) than DeepSeek V3 (5.6). (c) Number of distinct evaluation metric categories per candidate under three conditions (baseline, low-variety control, forced-depth intervention) for both models. Forced-depth raises this number substantially for both models; the low-variety control does not, confirming that the variety manipulation drives the effect. (d) Component-level score delta ($\Delta$, forced-depth – low-variety) for both models. DeepSeek V3 shows broad gains across all six rubric components; GPT-5 shows a large Approach gain but smaller or negative changes in other components.

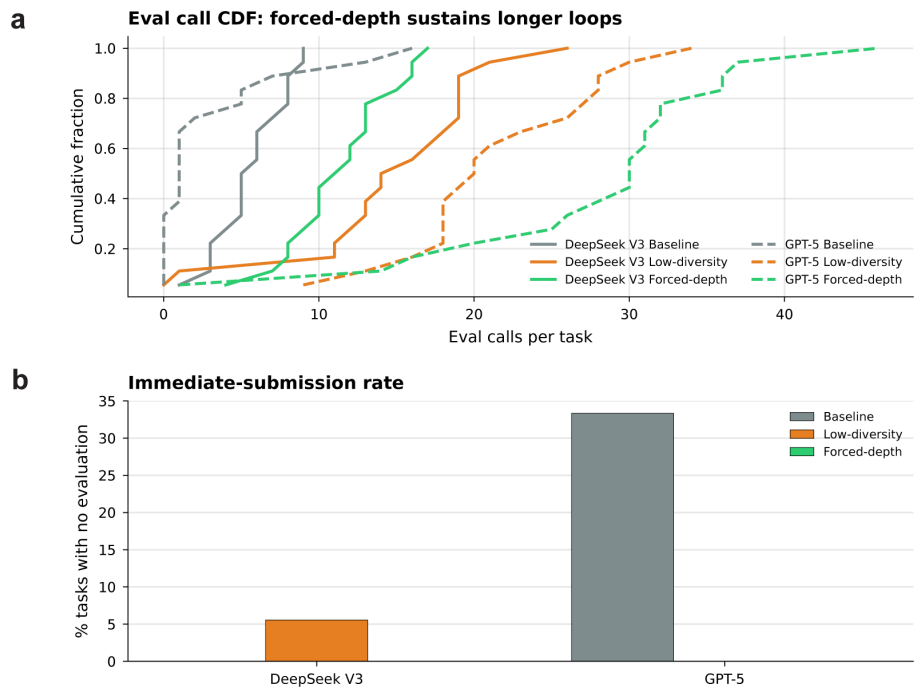

**Fig. S22 Early stopping and loop-termination behaviour under the intervention.** (a) Cumulative distribution function (CDF) of evaluation calls per task for DeepSeek V3 and GPT-5 under three conditions (baseline, low-variety, forced-depth). The forced-depth intervention shifts the CDF rightward for both models, indicating sustained longer evaluation loops. (b) Immediate-submission rate (percentage of tasks where the agent submitted a final answer with zero evaluation calls). GPT-5 baseline shows $\sim 33\%$ immediate submissions, which drop to 0% under forced-depth. DeepSeek V3 shows a low baseline rate ($\sim 5\%$ under low-variety) and 0% under forced-depth.

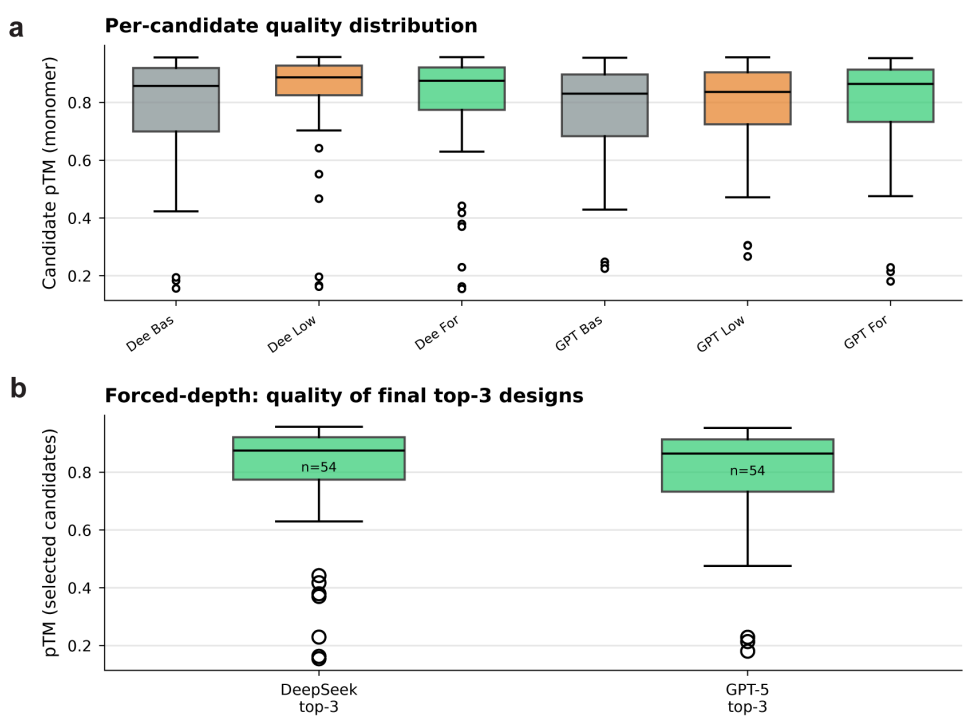

**Fig. S23 Candidate-level quality distributions and selection patterns under the intervention.** (a) Per-candidate pTM (monomer) distributions across three conditions (baseline, low-variety, forced-depth) for DeepSeek V3 (left three boxes) and GPT-5 (right three boxes). Candidate-pool quality distributions are broadly comparable across conditions, indicating that forced-depth does not alter the raw quality of generated candidates. (b) pTM distribution of the final top-3 selected designs under forced-depth for DeepSeek V3 ($n = 54$) and GPT-5 ($n = 54$). Selected designs show comparable median quality between models, confirming that the intervention improves selection rather than generation — agents retain higher-quality candidates from a similar underlying pool.

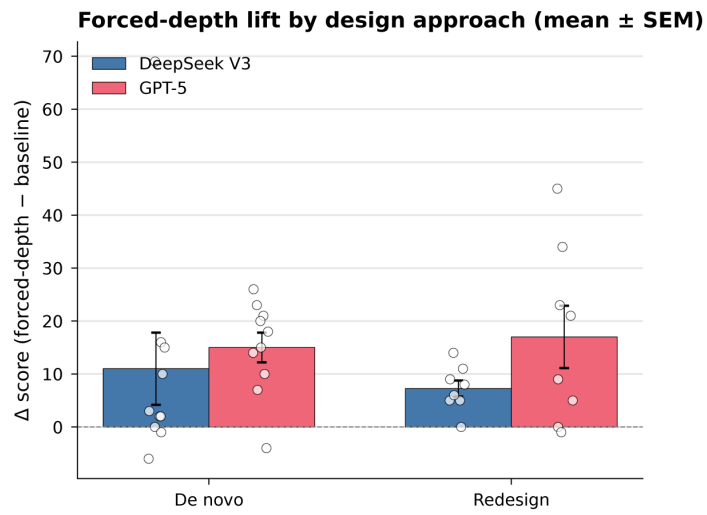

**Fig. S24 De novo versus redesign intervention effect.** Mean Δscore (forced-depth – baseline, ± SEM) split by design approach for DeepSeek V3 (blue) and GPT-5 (pink). Individual task deltas are shown as scatter points. Both models show positive intervention effects on both *de novo* and redesign tasks. The effect is modestly larger on *de novo* tasks for both models, mirroring the larger baseline depth-adaptation gap on generative tasks reported in Section M.

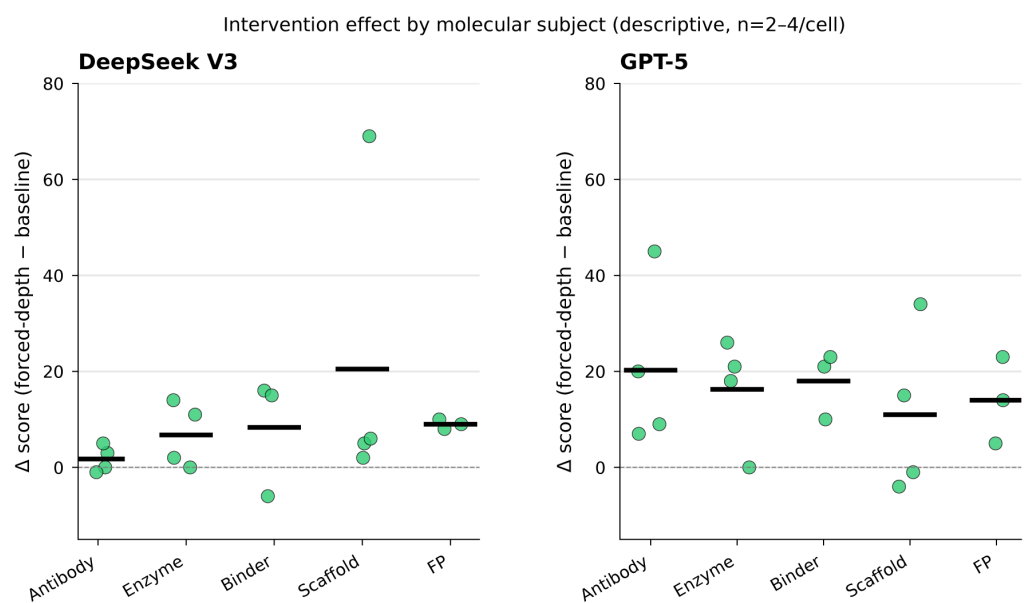

**Fig. S25 Domain-specific intervention heterogeneity.** Per-task $\Delta\text{score}$ (forced-depth – baseline) by molecular subject for DeepSeek V3 (left) and GPT-5 (right). Each dot represents a single task; horizontal bars indicate domain means. Due to the stratified subset design ($n = 2-4$ tasks per domain–model cell), these results are descriptive rather than inferential. Both models show positive effects across most domains, with the largest gains in scaffold and fluorescent-protein tasks for GPT-5.

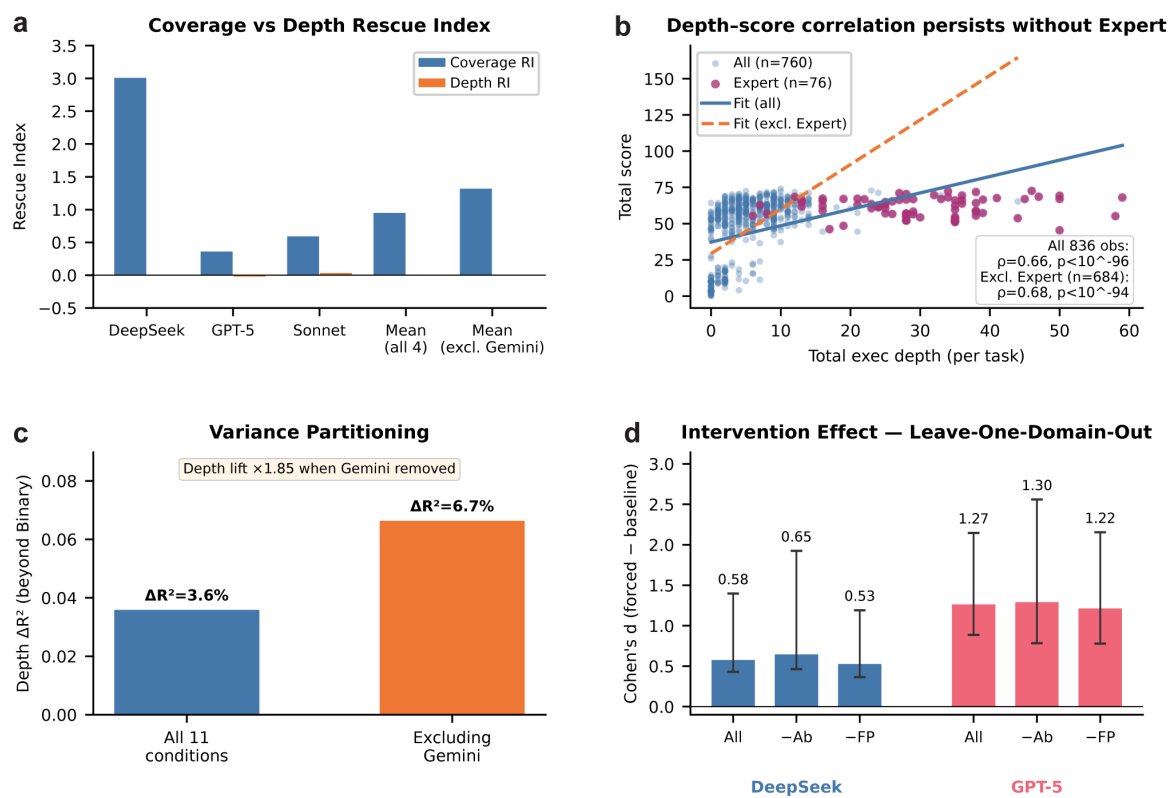

**Fig. S26 Robustness of central findings to Gemini exclusion and human-expert exclusion.** (a) Per-model unguided-to-guided coverage change versus depth change, computed with and without Gemini 2.5 Pro — the depth axis remains $\approx 0$ in both, confirming that the coverage–depth dissociation is not driven by Gemini’s tool-calling failure. (b) Variety of evaluation versus total score, with and without expert observations — the relationship persists ($\rho$ remains $> 0.6$), confirming that it is established across LLM conditions independent of the single-expert reference. (c) Hierarchical variance partitioning with and without Gemini — the variety axis’s explanatory contribution nearly doubles ($+3.6\% \rightarrow +6.7\%$) when Gemini’s floor effect is removed. (d) Forced-depth intervention effect sizes (Cohen’s $d$ with 95% bootstrap CI) under leave-one-domain-out resampling — effects remain large for both DeepSeek V3 and GPT-5 across all single-domain exclusions, confirming that intervention efficacy is not driven by any single molecular subject.
